# Estimating Historical Food Web Variation in Chesapeake Bay Using Isotope Variation in Museum Fish Specimens

**DOI:** 10.1101/2025.09.09.674953

**Authors:** Matthew P. Schumm, Katherine E. Bemis, Daniel K. Okamoto, Lynne R. Parenti

**Author notes:** Author contributions:* M.P.S. conceived the study with input from L.R.P., K.E.B., and D.K.O. All authors jointly collaborated on study design, selection of species and use of collections, and methods. M.P.S. processed specimens, collected samples, and performed data collection, with support from staff in the Division of Fishes, National Museum of Natural History, and the Museum Conservation Institute, Smithsonian Institution, and in the UC Davis Stable Isotope Facility. M.P.S. conducted analyses, made figures, and drafted the manuscript text with input from all authors.

## Abstract

Trophic interactions are hypothesized to change in response to rapid anthropogenic environmental changes. Documenting these trends may aid biological conservation. One tool to measure trophic level is stable isotope analysis. Nitrogen isotope values (δ^15^N) of bulk tissue may be used as indirect indicators of an organism’s trophic level and the isotopic composition of its prey. In contrast, compound-specific isotope analysis can estimate trophic level directly. For the fishes of Chesapeake Bay, museum collections provide a unique opportunity to characterize trophic trends collected as early as the 1850s, from an ecosystem that has been overfished and subjected to increasingly high nutrient loading and land use change during the past three centuries. To assess isotope data for evidence of change in predator species trophic level, we analyzed tissue from 183 museum specimens of three predators (Striped Bass, *Morone saxatilis*; Summer Flounder, *Paralichthys dentatus*; and Bluefish, *Pomatomus saltatrix*) and two lower-trophic-level species (Bay Anchovy, *Anchoa mitchilli*; and Menhaden, *Brevoortia tyrannus*). Predatory Striped Bass did not show an increase through time in δ^15^N values, despite such a trend in Bay Anchovy, suggesting possible change in trophic level. Analysis of Striped Bass tissue using compound specific analysis indicates their mean trophic level has been stable for decades. These findings imply that Striped Bass are part of a diverse food web within Chesapeake Bay beyond that reflected by our two included prey species and are stable through time in their average size-specific trophic level, a valuable insight into their ecological role.

## Introduction

Trophic interactions in aquatic ecosystems affect a broad range of phenomena from consumer populations’ dynamics (Buchheister et al. 2015) to the structure of entire ecosystems (Paine 1980). One characteristic of an organism’s trophic interactions are indices of trophic level, also referred to as trophic position, representing the mean level an organism occupies in a food web relative to the autotrophic primary producers at the base of the food web. Human fishing often initially targets higher-trophic-level fishes; thus highly fished communities may decline in maximum trophic level (Pauly et al. 1998; Essington et al. 2006; Branch et al. 2010). Yet even within a single species, variation in mean trophic level can result from a variety of factors. For example, fishing and other anthropogenic disturbances can lead to reduced body sizes within a species. This reduction may alter the prey species that the fishes target or are capable of capturing (e.g., Kindsvater and Palkovacs 2017). Moreover, trophic interactions may vary and change through time in fish of the same species and body size (Nakazawa et al. 2010) via changes in the mean trophic level of prey species and/or changes in the species composition of available prey.

Reconstructing past trophic levels may be challenging, but museum specimens provide one avenue to estimate shifts over time. Historical specimens can reveal a fish’s last meal via preserved gut contents (Beckmann et al. 2015). Preserved fish tissue also contains a record of trophic interactions in their stable isotope ratios that is more time-integrated than gut contents (Post 2002) – the relative abundances of common elements’ stable isotopes in preserved body tissues (Turner et al. 2023). These isotope ratios can be used to match predators to distinct prey sources (Ben-David and Flaherty 2012). Additionally, values of abundance of the isotope nitrogen-15 relative to the more common nitrogen-14, referred to as δ^15^N values, tend to increase in nitrogen-containing compounds as they are passed up trophic levels (Chikaraishi et al. 2009).

Knowing the length-specific trophic niche of a population informs ecosystem-based management (Engelhard et al. 2014). Isotope data can be pivotal to addressing such aims (Ólafsdóttir et al. 2021; Andrews et al. 2023). For example, data from the 1^st^ century CE to the present, from isotopes in bones of Atlantic Bluefin Tuna, *Thunnus thynnus*, collected from Mediterranean archeological sites, show a likely shift in tuna diets during the 16^th^-18^th^ Centuries. The diet shift was inferred primarily from stable isotope data for sulfur (δ^34^S values). The diet shift was possibly tied to habitat shifts in response to inshore anthropogenic exploitation of tuna prey (Andrews et al. 2023). Over a shorter timescale (forty years), Nakazawa et al. (2010) used δ^15^N values from a Japanese freshwater goby population to illustrate long periods of stability in length-specific diet punctuated by short-term change.

While isotopic analyses can provide a tool for reconstructing trophic niches, the method faces key limitations. δ^15^N values generally increase with trophic level (Post 2002). Yet primary producers can also vary in their δ^15^N values that propagate through the food web (Chikaraishi et al. 2009). “Bulk” δ^15^N measures nitrogen-15 relative abundance in all compounds within a tissue and is therefore affected both by a consumer’s diet and variation in δ^15^N derived bottom-up from primary producers (Chikaraishi et al. 2009) (Figure 1). To estimate trophic level from bulk isotopes, it is necessary to compare consumers of interest to primary producer δ^15^N values. For example, Willert et al. (2023) used museum fish collections and algae specimens from near Cape Cod. They found no increases through time in δ^15^N in two different fish species despite increasing δ^15^N in the algae, suggesting a decrease in these fishes’ length-specific trophic level. In contrast to bulk analysis, measuring δ^15^N values of specific amino acids, or compound-specific isotope analysis (CSIA), allows estimation of true trophic level (Chikaraishi et al. 2014). Some amino acids, including glutamic acid, are consistently affected in their δ^15^N values by the number of times molecules have been passed up the food web by consumption (Chikaraishi et al. 2009). Others, such as phenylalanine, are affected in this way at a rate of less than a tenth of that observed in glutamic acid (Bowes and Thorp 2015). The difference in trophic fractionation between glutamic acid and phenylalanine, i.e., the trophic discrimination factor for this pair of amino acids and their δ^15^N values, is well-studied for marine fishes (Bradley et al. 2015, see Figure 1 for a conceptual illustration). Comparing the δ^15^N values of amino acids therefore allows normalizing to the baseline δ^15^N values of producers, and estimation of trophic level from a single tissue sample. Moreover, fluid preservation of whole specimens does not affect trophic level estimates from CSIA (Durante et al. 2020; Welicky et al. 2021).

**Figure 1.**
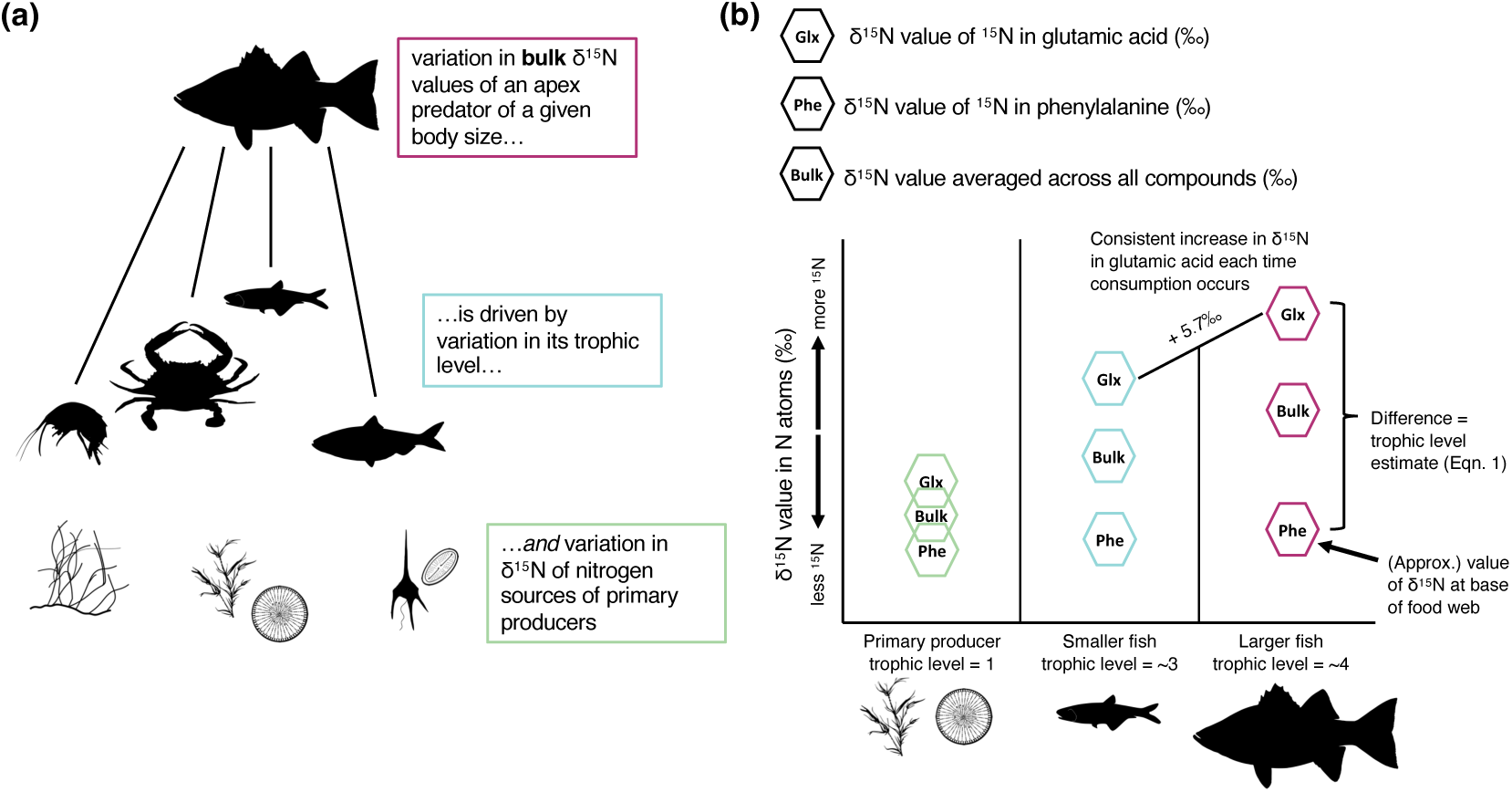
Conceptual figure of how δ^15^N values in body tissues change both due to trophic level, and due to variation in such enrichment existing at the base of the food web. Panel (a) shows a general illustration of this concept. The axes in panel (b) show how δ^15^N values in the nitrogen content of the amino acid glutamic acid consistently increase in fishes with additional trophic levels (transference of nutrients from a prey item to a predator), thereby also leading to a consistent increase with each trophic transference in “bulk” – for all nitrogen atoms in the tissue sample, across different compounds – δ^15^N values. In contrast, δ^15^N values in the amino acid phenylalanine remains close to the characteristic δ^15^N values at the nitrogen source for the base of the food web. Trends in the bulk δ^15^N values of tissues of apex predators may vary with bottom-up food web changes or changes in their own diet, whereas examining δ^15^N values at the level of specific compounds can be used to estimate trophic level from a tissue sample from a single organism.

Here, we used museum specimens collected over the past 170 years to evaluate the hypothesis that length-specific trophic level of predatory fish species in Chesapeake Bay has changed through time. More specifically, we used a combination of bulk tissue analysis of different fish species from muscle tissue, CSIA, radiography (X-ray) analysis of gut contents, and experimental estimation of specimen preservation bias on δ^15^N values. We hypothesized that bulk-tissue isotope ratios may have different temporal trends in predators as compared to prey. If spatial and temporal trends in δ^15^N values track closely between Chesapeake Bay predatory fish species and their prey species, this discrepancy would suggest that most change through time or space in the Chesapeake Bay food web is bottom-up and does not involve change in mean length-specific trophic level (Figure 1). If, for instance, δ^15^N values entering Chesapeake Bay food webs from primary producers were uniformly increasing, this increase would affect δ^15^N values of fishes throughout the food web. Change through time in predatory species trophic levels may be observed if temporal trends in δ^15^N values of their prey species are significantly different. For example, a decrease through time in body length-specific δ^15^N values of a predatory fish species, in a scenario where body length-specific δ^15^N values of their prey sources were increasing, would be expected if that predator was decreasing in its length-specific trophic level. We used a preservation experiment to quantify species-specific expected amounts of offset between observed δ^15^N values from preserved tissue, and values if measured from fresh tissue, due to the effects of preservatives themselves on δ^15^N values (e.g., Kishe-Machumu et al. 2017). We further hypothesized that, if temporal trends in bulk-tissue isotope ratios differed between prey and predator species, and if this indicated change in predator trophic level, then we would observe temporal change in isotope ratios in amino acids strongly fractionating with trophic level, in CSIA. If reflecting a temporal change in trophic level, this change would match that suggested by the bulk-isotope data – i.e., as in the earlier hypothetical example in this paragraph, if a predator were decreasing in its length-specific trophic level, we would observe a decrease in δ^15^N values of amino acids strongly fractionating with trophic level. We used CSIA on a subset of our samples from one species, Striped Bass, to further illuminate probable causes of patterns in their isotopic variation, and whether trophic level in their population has changed over time.

## Methods

### Study system

The Chesapeake Bay has been heavily affected by human harvest and other impacts especially since European colonization (Black et al. 2017), leading to changes in trophic dynamics (Watts et al. 2024). For some taxa there is strong evidence for changes in length-specific diet over time since the 1950s, especially due to some forage fishes (chiefly Menhaden) being heavily harvested (Overton et al. 2015). These potential shifts in trophic dynamics are complicated by spatial heterogeneity in habitat’s biological and physical structures, and in availability of particular prey items (Overton et al. 2009). Many fish species spend much of their lives not just within Chesapeake Bay, but in multiple, distinct river systems that radiate out from Chesapeake Bay. These river systems have different environmentally available δ^15^N values (Davias et al. 2014). Moreover, there have been substantial changes in anthropogenic nutrient inputs to Chesapeake Bay (Pan et al. 2021). These inputs potentially affect not only bottom-up productivity, but also δ^15^N values that are signatures of primary producers that may be transmitted to herbivores and predators. δ^15^N values percolating up Chesapeake Bay food webs have changed because they tend to be higher in anthropogenic nitrogen input (Fertig et al. 2014). Specifically, anthropogenic impacts including nutrient loading and associated hypoxia have altered food webs (Long and Seitz 2008) and human impacts have removed habitat types such as oyster beds and wetlands that previously slowed and buffered nutrient flows (Kemp et al. 2005).

We took measurements and tissue samples from individuals from each of five Chesapeake Bay fishes (Striped Bass, *Morone saxatilis*; Summer Flounder, *Paralichthys dentatus*; Bluefish, *Pomatomus saltatrix*; Bay Anchovy, *Anchoa mitchilli*; and Menhaden*, Brevoortia tyrannus*). These species are abundant and important components of the Chesapeake Bay ecological community (Buchheister and Latour 2015). We selected these species because there are extensive data on current and recent diets and other biological characteristics (Pruell et al. 2003; Walter and Austin 2003; Overton et al. 2009; Szczebak and Taylor 2011; Buchheister et al. 2015; Overton et al. 2015), including data on their populations in Chesapeake Bay. We chose species with abundant information on their diet with which we could interpret our findings. Striped Bass, Summer Flounder, and Bluefish are all predators of fishes and invertebrates; Bay Anchovy is an opportunistic zooplanktivore and detritivore, and Menhaden is a filter-feeder consuming both phytoplankton and zooplankton (Collette and Klein-MacPhee 2002).

### Selection of taxa and specimens for bulk isotope analysis

We used 196 preserved and accessioned specimens at the Smithsonian Institution National Museum of Natural History (NMNH) Fish Division and the Nunnally Ichthyology Collection at the Virginia Institute of Marine Science (VIMS) (see “Specimens Examined”, Sheet 1 of the Supplemental Data Table). Specimens of the two smaller and lower trophic level species, Bay Anchovy and Menhaden, were available in the NMNH and VIMS collections in larger numbers than specimens of the other three taxa. For Bay Anchovy and Menhaden, we divided the available specimen *lots* (all specimens of one species collected at one time and stored in a single jar) into different combinations of covariates to balance our sampling across spatial and temporal contexts. We divided available specimen lots into (i) quarter-century time periods between 1900 and 2000, (ii) seasons (whether they had been collected within or outside of April-September), (iii) regions of the Chesapeake Bay corresponding to different river watersheds flowing into the Bay, and (iv) whether or not they had white eye lenses (generally indicative of initial fixation in alcohol rather than formalin, De Bruyn et al. 2011). Then we selected about 40 individuals representing approximately equal numbers, where possible, of specimens from each combination of variables.

For the three larger and higher trophic level species, Striped Bass, Summer Flounder, and Bluefish, fewer specimens were available. Smaller species are often easier to capture and more economical to store in large numbers in museum collections (Gotelli et al. 2023). Thus, for these three taxa, we sampled from all individual fish specimens of 15 cm standard length (SL; from tip of the snout to distal end of the caudal peduncle) or longer in the NMNH fishes collected from Chesapeake Bay. At VIMS, we also sampled from all 15 cm SL or longer Bluefish and Striped Bass from Chesapeake Bay. We sampled two Summer Flounder (selecting randomly the two lots to use, and the individual specimen within the lot to sample from) from the 1960s, a decade not represented in the NMNH collections for Summer Flounder. Figure 2 shows the distribution of specimens we sampled across different space and time categories that were retained in our statistical analysis. Supplemental Table 1 summarizes this information as well as for additional covariate categories. Supplemental Figure 1 is a map of the sites from which Striped Bass in this study were collected. Only 183 of the 222 specimens (both previously preserved fishes, and fresh fishes described later in the Methods) that we sampled were retained in our final statistical analysis for bulk tissue, because some specimen lots lacked environmental covariate data, even following a review of hand-written jar labels, field notes, and ledgers, that would inform our statistical analyses.

**Figure 2.**
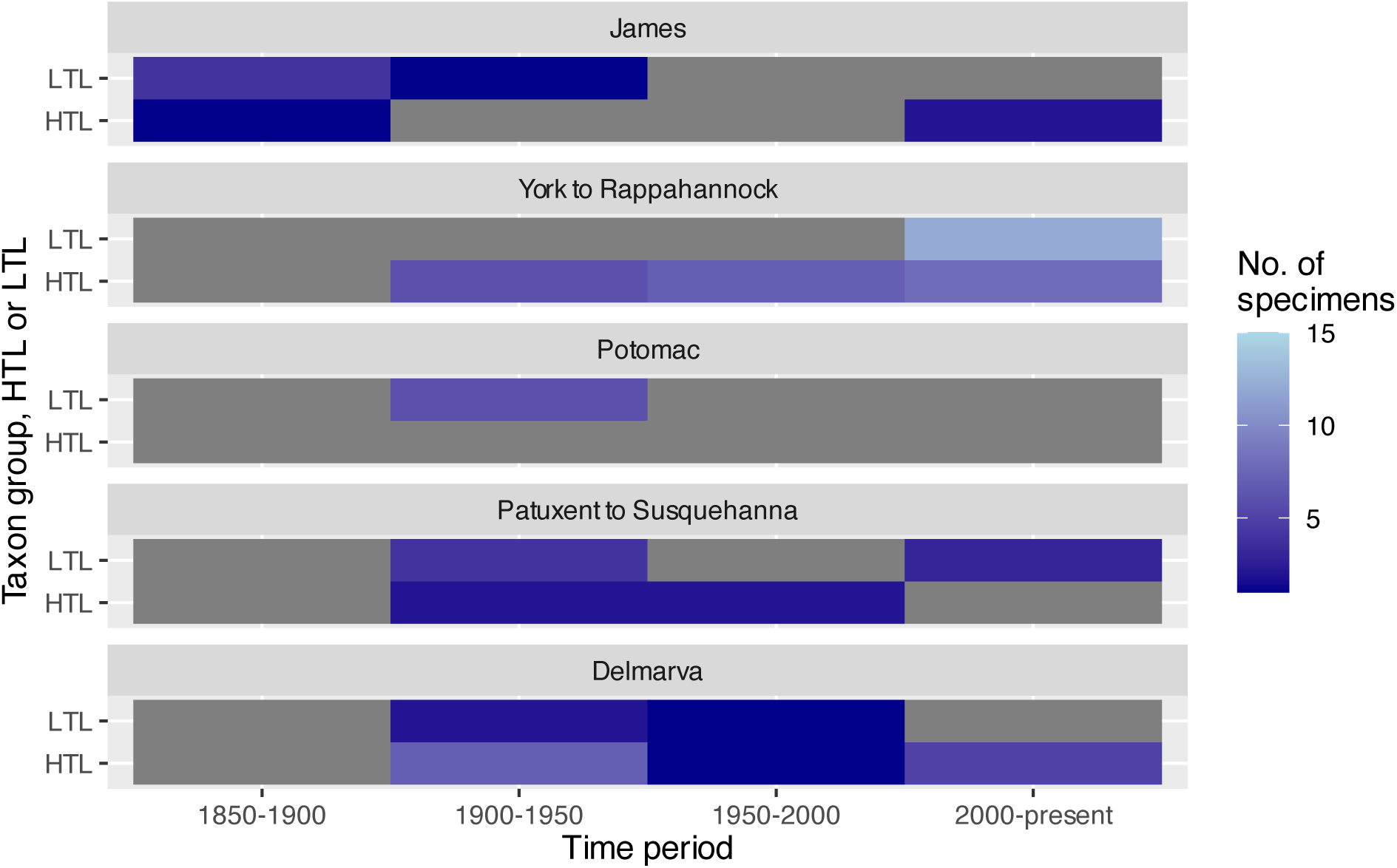
The number of total specimens from which we collected and analyzed bulk δ^15^N values from each of the combinations shown in this figure of group of species, time period, and region of the Chesapeake Bay. “HTL” on the y-axis for taxon group denote species known to generally be of higher trophic level out of the five species we studied – Bluefish, Striped Bass, and Summer Flounder, and “LTL” denotes lower trophic level species, Bay Anchovy and Menhaden.

Our goal in exploring evidence for possible trophic level shifts in predatory Chesapeake Bay fishes was to test for change in trophic level among individuals of a given length. Therefore, for each of our three higher-trophic-level species, we focused on fishes between 15-30 cm in standard length (SL), a small subset of these species growth trajectories. At standard length (SL) of ≥ 15 cm, niche shifts occur in Striped Bass, Summer Flounder, and Bluefish and therefore ≤ 15 cm individuals may be part of different, incomparable food webs (Walter and Austin 2003; Szczebak and Taylor 2011; Cernadas-Martín et al. 2021). Thus, for these three species, we generally avoided sampling from individuals under 15 cm. However, 15.0% (22 out of 147) of our sampled Striped Bass, Summer Flounder, and Bluefish individuals are between 9-15 cm and were sampled when they had a covariate combination (e.g., combination of watershed and decade) not represented in our data in ≥ 15 cm fish, and/or where they were in the same jar as other larger specimens. Having specimens lots for which we sampled multiple preserved individuals increased our ability to accurately estimate random effects for lot-specific effects in our mixed-effects statistical models and therefore to better estimate fixed effects for time and other covariates. Focusing on specimens of these species at a larger range of standard lengths when they would be more targeted by fisheries was not an option because of how few large-bodied fish specimens are present in museum collections. Fish specimens larger than ∼40 cm were not present in the collections for these species. This limitation of museum fish collections means our sampled specimens likely include Bluefish no older than 2 years, Striped Bass older than young-of-year but no older than 2-3 years, and Summer Flounder older than young-of-year but no older than 5 years (Collette and Klein-MacPhee 2002; Latour et al. 2012).

We also included fishes in this study that were not already preserved in museum collections, primarily to study δ^15^N values before and after preservation but also to add to the spatiotemporal breadth of our isotope data. We requested specimens of these five fishes from the survey sampling of the Chesapeake Bay by VIMS in their Chesapeake Bay Multispecies Monitoring and Assessment Program (ChesMMAP) and Juvenile Finfish Trawl Survey, from their collection efforts in late 2021 and 2022. These fishes were to be used for an experiment studying effects of preservatives on measured δ^15^N values and we incorporated the bulk-tissue δ^15^N values from these fishes into our dataset with a covariate in our analyses to adjust for their being freshly caught and unpreserved. These fishes are also included in Supplemental Table 1 and Figure 2. Virginia state law (the Code of Virginia, section 28.2-1101) authorizes VIMS to harvest fish from state waters for research purposes, serving as the collection permit for that work, and these specimens were deeded as a gift by VIMS to NMNH. These fresh fishes were then catalogued at NMNH (see “Specimens Examined”, Sheet 1 of the Supplemental Data Table). The isotope data taken from them before fixation was included with preserved tissue samples in our full data analysis for bulk tissue isotopes, with covariates for preservation to account for these specimens being fresh.

For each fish specimen, we measured the standard length (SL) and weighed it after patting off excess preservative fluid with paper towels, following best practices for collecting length-weight data from museum fluid-preserved specimens (Hay et al. 2020). We used a 6 mm diameter biopsy punch tool to take 50-100 mg wet dorsal muscle tissue from the right side of the fish just anterior to the dorsal fin(s). We checked for ethanol fixation indicated by bright white preserved lenses. We also recorded (when possible) all collection details including location and date of collection. All specimens we sampled were known to be from Chesapeake Bay and/or the Delmarva Peninsula. For our final data analyses, we only retained specimens with associated data on year and month of collection. For samples of muscle tissue for analysis of δ^15^N values as a proxy for diet variation, tissue samples are often processed to remove their lipid content prior to further laboratory analysis. This process is justified because lipid concentration may affect measured δ^15^N values and variation in lipid content between samples may add variation in δ^15^N values that is not due to true diet variation (Sotiropoulos et al. 2004). Lipids generally have been removed from the subcutaneous muscle tissue of fluid-preserved fish specimens by the process of formalin and ethanol fixation and preservation (Edwards et al. 2002). Isotope analysis on muscle tissue from fluid-preserved museum fish specimens typically has not included a lipid extraction step (Turner et al. 2015; Durante et al. 2020; Welicky et al. 2021).

### Experimental preservation to assess preservative bias in isotope data

We used multiple approaches to accounting for biasing effects of museum specimen preservation on δ^15^N data in our study. Museum fish specimens typically are fixed using a formalin solution, although some older specimens have not been fixed in this way, and they are then stored and preserved long-term in ethanol solution (De Bruyn et al. 2011), both of which may affect δ^15^N values (Durante et al. 2020). Despite such effects, isotopic methods may still provide useful information about an individual’s diet prior to their death and preservation. The effect of preservation on bulk tissue δ^15^N values is expected to occur within less than a year of fixation rather than slowly over time, and to be constant for a given species, preservation method, or lipid content at the time of preservation (Durante et al. 2020). For example, Kishe-Machumu et al. (2017) found for Lake Victoria cichlid specimens that this effect is small relative to patterns of natural temporal variation in fish tissue δ^15^N values. Moreover, fixation does not affect the estimates of trophic level that can be made by compound-specific isotope analysis (Durante et al. 2020; Welicky et al. 2021). Thus, if we account for short-term bias imposed by preservation methodology, isotope analysis can be a potentially powerful tool for analyzing shifts in trophic level over time using museum specimens. We focused primarily on temporal variation in δ^15^N values in specimens that had been preserved and in collections for several years. Consequently, we assumed that the influence of preservation on the δ^15^N values of a fish of a given species was a constant value across time periods. We included fresh specimens in our dataset and statistical analyses. We estimated and fit a constant effect of preservation for the preserved specimens as compared to fresh specimens. Fitting an effect for preservation on δ^15^N values allows fresh and preserved fishes to be compared within the same statistical model. We also fit an effect in some of our models for the difference in δ^15^N values between fishes specifically fixed with formalin vs. those without it.

We also conducted an experiment sampling fresh fishes before and after preservation procedures. This experiment was to further validate our assumptions about whether preservation effects occur quickly upon a specimen relative to the historical time range of the museum specimens we studied. Other studies had shown, with different species than in our study, that such preservation-caused biases occur soon after preservation and do not increase over time as indicated by other studies (Kishe-Machumu et al. 2017; Durante et al. 2020). We also aimed to estimate the magnitude of such bias, to contextualize our inferences. Specifically, we aimed to estimate the magnitude of preservation-induced increases in δ^15^N relative to other effects. For this purpose, we used 15 fresh (frozen) fishes (3 from each species) to measure change in isotope ratios (i) after formalin fixation with a 10% solution for one week and (ii) after preservation in 75% ethanol, both shortly after preservation and after over a year of preservation. We sought to test for a statistically significant increase in δ^15^N values following typical preservation methods for these particular species (for which this bias had not been previously measured), and to further observe if there was any statistically significant difference between the preservation-biased δ^15^N values (as contrasted with fresh tissue values) when measuring after over a year had passed as opposed to measuring δ^15^N values within a month of initial preservation. No significant change in the effects of preservation in over a year and a half would be consistent with preservation biases not being confounded with time, which is an assumption we make in our analyses. We also used 11 individuals from these surveys (2 specimens of Menhaden, none of Striped Bass because of limited availability of these species, and 3 specimens of each of the other species) to measure effects of ethanol preservation without prior formalin fixation. Isotopic data was obtained from these specimens before fixation and preservation to compare with post-fixation and post-preservation.

### Bulk isotope laboratory analysis

Tissue samples taken from fluid-preserved fishes were stored in cryovials and 1.5ml plastic sample tubes in a −20°C freezer prior to bulk isotope analysis. Prior to preparation and packing in tin caps for isotope analysis, all tissue plugs were dried in a drying oven at 60°C for at least 24 hours. Dried tissue plugs were ground inside their sample tubes with a plastic pestle, or with a mini-mortar and pestle. Tools were wiped down with acetone between different samples. Dried samples crushed to a fine powder were then packed in tin caps folded into a box shape and put in sample trays to be loaded into the isotope ratio mass spectrometer at the Stable Isotope Facility at the Smithsonian Institution Museum Conservation Institute. All samples were run on a Thermo Delta V Advantage isotope ratio mass spectrometer in continuous flow mode coupled to an Elementar vario ISOTOPE Cube Elemental Analyzer (EA) via a Thermo Conflo IV. All calculations of raw isotope values were performed with Isodat 3.0 software. A set of reference materials was included for every 10-12 samples. All reference materials were run with the same parameters and procedures as samples, and isotope values were corrected using a linear correction on the calibrated reference materials. The error associated with all sample data points was <0.2%. This estimate of reproducibility was based on repeated spectrometry of selected reference and sample materials. The isotope values were reported in units of per thousand (‰) and were calculated relative to atmospheric air as a standard, where the δ value for an isotope (here, ^15^N) is calculated as [R_sample_-R_air_]-1. R represents the ratio of the rare stable isotope to the more common isotope of the element (here, this ratio is ^15^N/^14^N). A higher δ value therefore corresponds to a higher ratio of ^15^N/^14^N, as compared to the standard of the atmosphere’s gaseous nitrogen. The δ^15^N value of the atmosphere’s gaseous nitrogen serves as a standard for stable isotope laboratory analyses.

### Statistical analysis of bulk isotope data

We modeled δ^15^N value measured from the muscle tissue of an individual fish as a function of fish standard length, region or watershed in which the fish was collected, year of collection (as a continuous variable), whether the fish was preserved or fresh and frozen, whether it was from a specimen that had been formalin-fixed prior to tissue sampling, and its species. We allowed Year interactions with Species and with the River/Watershed variable. We fit the relationship of our δ^15^N value outcome to these explanatory variables using a Bayesian generalized linear mixed-effects model using the R package brms (Bürkner 2017). We accounted for nonindependence by including random effects for fish from shared specimen lots. We used the package bayestestR (Makowski et al. 2019) to calculate Bayesian probabilities of effects in a certain direction (effect sizes above or below zero). We included standard length as a covariate in our model to standardize our assessment of potential temporal trends in δ^15^N values in bulk tissue to fish of a given body size.

We compiled data from the literature to include as covariates in our analysis on environmental variables relevant to explaining spatiotemporal variation in δ^15^N values, but we did not include all of these variables in the model we interpret in and describe in detail in the Results. We opted in the final model version to accommodate environmental differences fitting mostly via just one environmental variable (the watershed from which the fish was collected), to let the random effects for individual specimen lots fit and absorb other between-specimens variation, and to exclude some additional environmental variables (Season and Salinity) that were not significant predictors of δ^15^N in our analyses. The different watersheds emptying into Chesapeake Bay differ in many environmental variables, including how they are affected in nutrient flows by anthropogenic activity (Pan et al. 2021). This environmental heterogeneity led us to include the river watershed the specimen was collected from or nearest to as a covariate. We grouped fishes found in and around the Delmarva Peninsula’s Chesapeake Bay and Atlantic coasts, as opposed to those found in or near one of the rivers on the Chesapeake Bay western shore, in a single category for this variable. Grouping all Delmarva specimens, and grouping all of the western shore specimens from the Upper Chesapeake Bay (including those from near the Susquehanna) as “Upper Chesapeake Bay - West” specimens, resulted in specimens from at least four of our five study species for each of the seven levels of this covariate: the James, the York, the Rappahannock, the Potomac, the Patuxent, Upper Chesapeake Bay - West, and Delmarva.

Additionally, salinity varies geographically within Chesapeake Bay, and varies seasonally, with both salinity and season often having effects on nutrient flows that can significantly contribute to stable isotope variation in estuarine fishes (Suzuki et al. 2008). Consequently, we used maps of mean surface salinity in Chesapeake Bay as a coarse estimate of salinity at the place and time a fish was collected (USEPA 2019). We did not use season-specific salinity values for this scoring so that variation in this salinity covariate would not be confounded with variation in our Season covariate. We also included in our dataset whether the fish was collected in Winter or between the Fall and Spring equinoxes (i.e., collections from October through March were counted as Winter) as a binary Season covariate.

We chose the final version of the model to focus on primarily in our Results by increasing the complexity and flexibility of model fitting methods while discarding model variables without significant or substantial contribution to variation in our data and focusing on model terms with an intuitive interpretation supported by exploratory analysis of the data.

Stochastic, Bayesian model fitting procedures for generalized linear mixed-effects models allow incorporating uncertainty from fixed and random effects without computing complex marginal likelihoods (Bolker 2015). However, our conclusions were robust to model fitting techniques. We fit a generalized linear model (trying versions both with and without random effects) with a frequentist package lme4 (Bates et al. 2015), removed insignificant environmental covariates as determined by F-test comparisons of models, and then fit a Bayesian model that did not qualitatively differ from the frequentist models in magnitude and direction of covariates. We also removed body weight and its interaction with standard length from the model as these variables were not significant. Priors used for Bayesian models were Normal distributions with standard deviation of 10. The results of analyses run with these priors did not differ qualitatively from the effects of using default brms priors. A version of the Bayesian model with a Gamma likelihood and log-log power law relationships allowed between δ^15^N value and our continuous variables (standard length, year) had a significantly worse (higher) WAIC score than a model with simply a Gaussian likelihood and identity link function. A version of the Bayesian model with nonparametric nonlinear splines or smoothers for (River Watershed * Year) and (Taxon * Year) relationships through time is described briefly in the Results but also had a worse (higher) WAIC score than a model with simply a Gaussian likelihood.

### Radiography of gut contents

Using a DURASCAN 1417 digital x-ray system, we radiographed specimens of higher-trophic-level fishes to examine indications of their diet just prior to collection. We radiographed all of the specimens of the three higher-trophic-level species that we sampled for tissue. The x-rays for this study were taken at exposures of 40 kilovolts for 10 seconds. Raw X-ray files were converted to image files using 16-bit Signed format in ImageJ, with black as zero. Fishes were scored as with or without vertebrate prey in their guts, a proxy for piscivory.

We statistically modeled the outcome of whether we observed vertebrate prey in specimens’ guts as a Bernoulli distributed random variable taking values of either 0 (no vertebrate prey) or 1, using a generalized linear mixed-effects model fit in brms as described in the section of the Methods on statistical analyses of the bulk isotope data. We used standard length of the fish and year as predictors.

### Compound specific isotope analyses to directly estimate variation in trophic level

We examined our bulk isotope results for evidence of predators showing a statistically significantly different temporal trend in bulk muscle tissue δ^15^N values relative to the temporal trends in common prey items. Such difference in temporal trends might indicate that a predator varied through time in its mean trophic level relative to this prey item, as opposed to variation in its δ^15^N values simply due to variation in δ^15^N values at the food web source. Based on bulk isotope results, we identified Striped Bass as possibly showing trends in δ^15^N values reflecting a particular factor, such as a change in trophic level, rather than just changing δ^15^N values at the source of Chesapeake Bay food webs.

Change in trophic level cannot be confirmed from examining bulk isotope results of a predator and one of its prey items, but compound specific measurements of δ^15^N values can normalize δ^15^N values for accurate measurement of trophic level (Chikaraishi et al. 2009). Following similar work (Welicky et al. 2023), we sent Striped Bass samples for compound-specific amino acid nitrogen isotope analysis to the UC Davis Stable Isotope Facility. These compound-specific isotope measurements allow separation of trends in isotope variation caused by variation at the base of the food web from those caused by increase in δ^15^N values as energy passes up the food web. We used the δ^15^N values for phenylalanine and glutamic acid because glutamic acid increases in δ^15^N values as food is transferred up trophic levels by a known mechanism, whereas phenylalanine increases in δ^15^N values with increasing trophic level at less than a tenth of the rate observed in glutamic acid. Further, a formula exists to estimate mean trophic level from these amino acids (Bradley et al. 2015, see Figure 1 for a conceptual illustration), and this estimate is robust to museum preservatives (Welicky et al. 2021). The estimated mean trophic level for a sample analyzed for compound-specific nitrogen-15 of nitrogen atoms is calculated as:

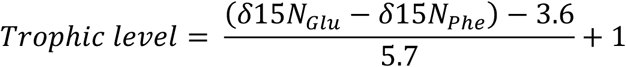

where 3.6 is the difference (in units of ‰ ^15^N/^14^N compared to the standard of nitrogen gas in air) that exists between δ^15^N values for glutamic acid and phenylalanine even at the base of the food web, 5.7 scales the resulting difference to units of trophic levels, and the addition of 1 accounts for the fact that primary producers (the bottom of the food web) are defined as 1 on the scale of trophic level. The term trophic ‘position’ is sometimes used for such a continuous value ranging from 1 upward, with trophic ‘levels’ referring to discrete groupings of species at distinct values of trophic position (e.g., Thompson et al. 2007). Here, we use ‘trophic level’, and intend it to be interchangeable with ‘trophic position’, as in Pasquaud et al. (2010). Higher trophic levels or positions, with values above 1, then represent continuous-valued averages of the number of times an organism’s organic molecules have passed up the food web. For example, an organism with a diet of roughly 50% primary producers and 50% primary consumers might have a trophic level of 2.5.

For compound-specific stable isotope analysis, we used remaining powdered tissue from Striped Bass samples taken for bulk isotope analysis as described in the Methods. We used Qorpak Clear Glass Threaded Vials with PTFE Lined Caps to ship samples to UC Davis Stable Isotope Facility. The lab used acid hydrolysis to remove amino acids from sample material proteins (6 M HCl, 70 min, 150 °C under N_2_ headspace). Amino acids were then derivatized as *N*-acetyl methyl esters, and derivatives were separated on an Agilent DB 35 column (60 m X 0.32 mm ID, 1.5 µm film thickness). Following this step, gas chromatography was performed using a Thermo Trace GC 1310 gas chromatograph coupled to a Thermo Scientific Delta V Advantage isotope-ratio mass spectrometer via a GC IsoLink II combustion interface. The acceptable measurement error was <1%, as assessed by repeated measures of reference materials (quality control and assessment materials were measured every 5 samples in the stable isotope mass spectrometry run).

For statistical analyses of these data, we modeled the estimated trophic level, calculated from the measured glutamic acid and phenylalanine δ^15^N values, as a function of fish standard length and year. We used a similar model structure to the main preferred statistical model we report on in our Results for our bulk analysis, with the outcome modeled as Gaussian with an identity link to the predictors, as described in the Methods. We also ran a version of this model including covariates for Season (Spring, Summer, Fall, and Winter, using the astronomical definitions).

## Results

### Experimental preservation: measuring effects of preservatives on specimen isotope chemistry

Effects of preservation from different museum fixatives appear small and fast acting, compared to effects of other covariates, as measured by experimental preservation of freshly collected and frozen specimens. Effects occur and stabilize within weeks, in comparison to over years or decades, which would confound conclusions made about temporal change. 10% formalin fixation of 15 fresh specimens for seven days followed by transfer through graded series of alcohol to 75% ethanol increases δ^15^N values relative to fresh tissue by 0.83‰ (SE = 0.20) compared to fresh and not fixed samples (Figure 3). Similarly, 11 fresh fish samples immersed in 75% ethanol for 14 days show an increase in δ^15^N values by on average 0.82‰ (SE = 0.09) compared to fresh and not fixed samples (Figure 3). After 19 months of preservation, we revisited these specimens and found no additional statistically significant change in their δ^15^N values (Figure 3).

**Figure 3.**
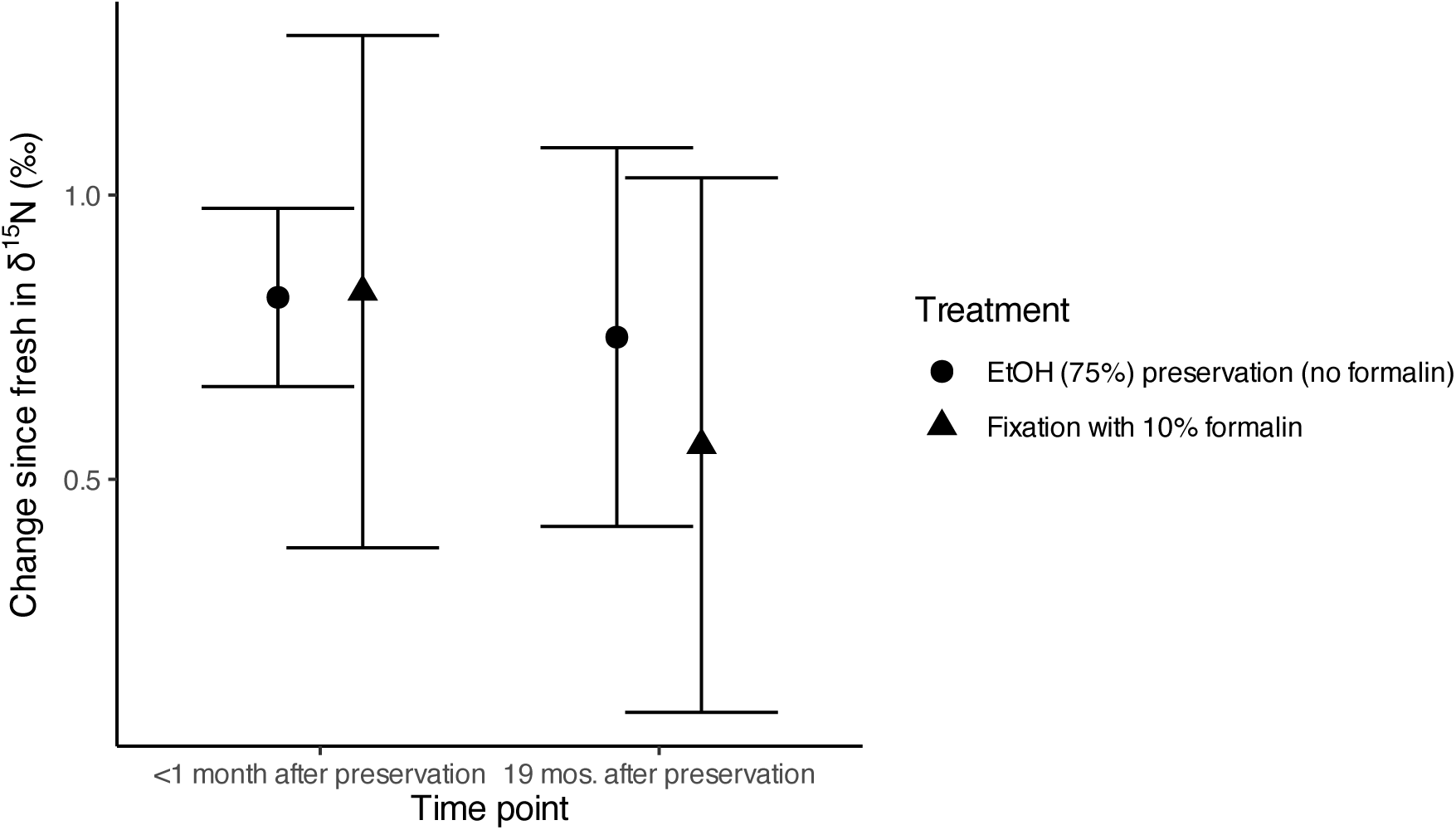
Effects of preservation, at two time points post-preservation, on the δ^15^N values for a dorsal muscle tissue plug taken from a fish specimen. Error bars are +/- 1.96 times the standard error of the values for the change in a specimen’s δ^15^N values.

### Bulk isotope analysis: temporal trends in bulk δ^15^N

Our bulk isotope data show increasing trends of δ^15^N values over time across all specimens taken together, with a significant difference between the trend through time in δ^15^N values for Striped Bass as compared to that for their common prey item, Bay Anchovy. These trends were fitted by our statistical model while accounting for other covariates (Table 1). The raw data for bulk δ^15^N values are shown in Figure 4a-e, with simple δ^15^N by time smoothers evaluated at two points – without other covariates included – fit to the data. As detailed in Table 1, the slope of the estimated trend in δ^15^N varied among species. The trend is largest (most positive) in Bay Anchovy compared to the other four species. For Striped Bass, the slope of change in its δ^15^N values for muscle tissue is shallower than that of the Bay Anchovy (95% credible interval of the difference = −1.92 to −0.14, Table 1) from the 1880s to 2020s. Menhaden and Bluefish also show a significantly different time trend from Bay Anchovy in Table 1, whereas Summer Flounder does not. The estimated probability that the change in Striped Bass muscle tissue δ^15^N values has a slope with time that is either negative, or not as strongly positive, as that for Bay Anchovy, is 98.38%, based on the Bayesian posterior probability density. Bay Anchovy estimated δ^15^N values increased on average from the 1880s to 2020s. For a median-standard-length Bay Anchovy (6 cm SL), the estimated δ^15^N value changes from 14.3 in the 1880s to 17.8‰ in the 2020s, compared to a change from 16.8‰ to 17.5‰ for a 22 cm SL Striped Bass (Figure 5a). The general trends of increase, and the disparity between Striped Bass and both Bay Anchovy and the other three taxa, are apparent qualitatively from observing the raw data points in Figure 4. This difference in the raw data is consistent with Striped Bass having the slope for δ^15^N values with time that differed the most from Bay Anchovy’s slope in Table 1’s statistical analysis, even compared to the other three taxa and how much they differed from Bay Anchovy in Table 1. Supplemental Figure 2 shows the nonlinear nonparametric trends that were fit by a model using splines for taxon-by-time relationships. Including these splines resulted in a worse (higher) WAIC score for the statistical model, and do not qualitatively suggest that there are nonlinear trends or differences between species that were not captured with our linear models.

**Figure 4.**
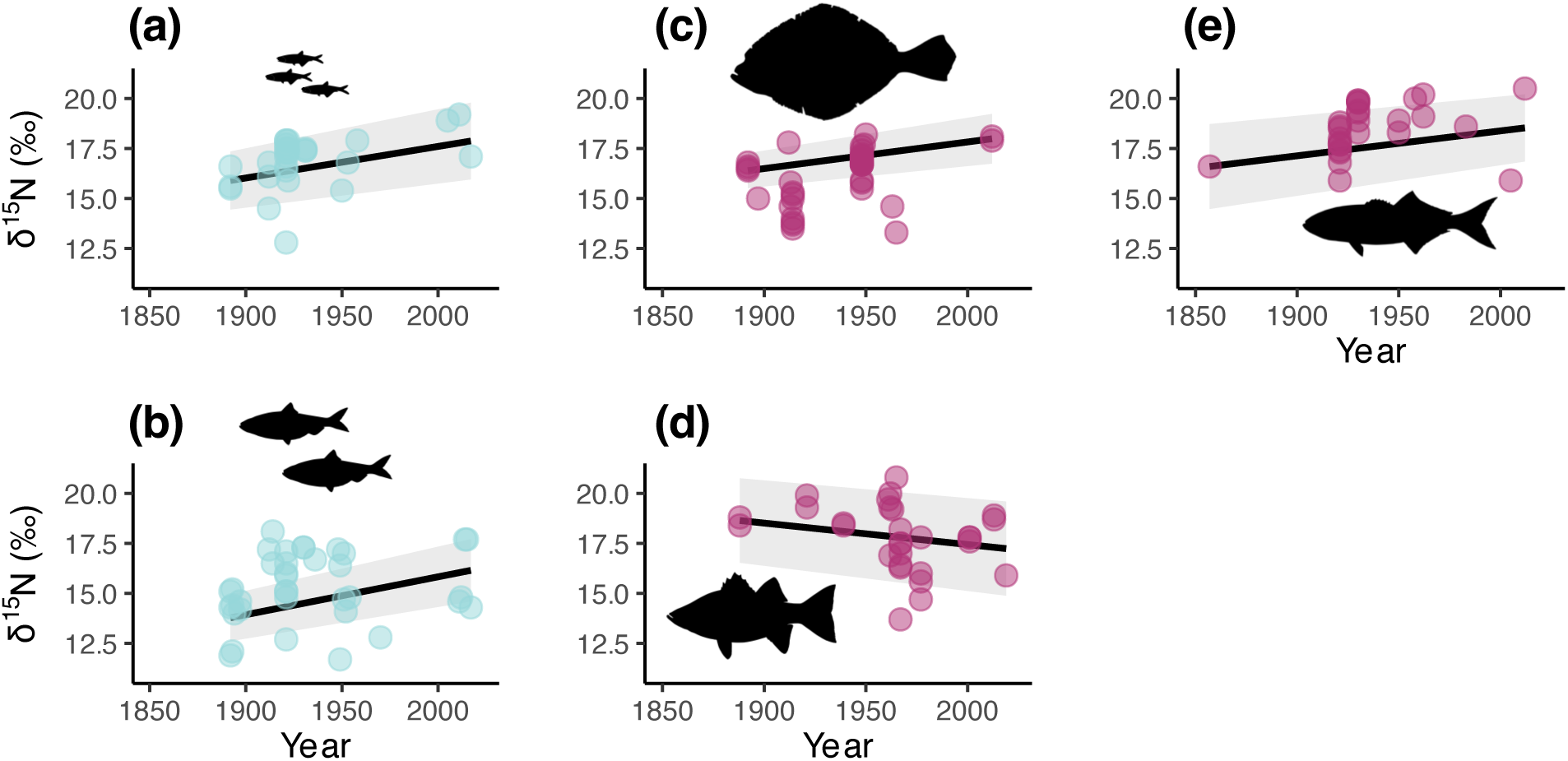
(a-e) Bulk (for all nitrogen atoms in the tissue sample, across different compounds) δ^15^N values (relative to amount of the nitrogen-15 isotope in atoms in N_2_ gas in air) for 157 preserved museum specimen tissue samples from five sampled Chesapeake Bay fish species. (a) Bay Anchovy, (b) Menhaden, (c) Summer Flounder, (d) Striped Bass, and (e) Bluefish. X-axes show the year of specimen collection. Individual points are raw data. The fitted line and confidence interval are for an exploratory LOESS smoother evaluated at two points to show the general linear trend in the data prior to addition of other explanatory covariates. Fishes from 2021-2022 that were collected fresh and included as part of our isotopic dataset were excluded from these plots. Isotope values from the fresh specimens may vary substantially from other data points because they are from unpreserved tissue, which is an effect that is parsed out in our tables by our statistical modeling of these data. These plots are intended to show the general trend with time of δ^15^N before taking into account effects of other covariates such as preservation method of the sampled fish, hence our exclusion of the fresh specimens from these plots (which only represent the near present, and thus are not visually informative as to temporal trends).

**Figure 5.**
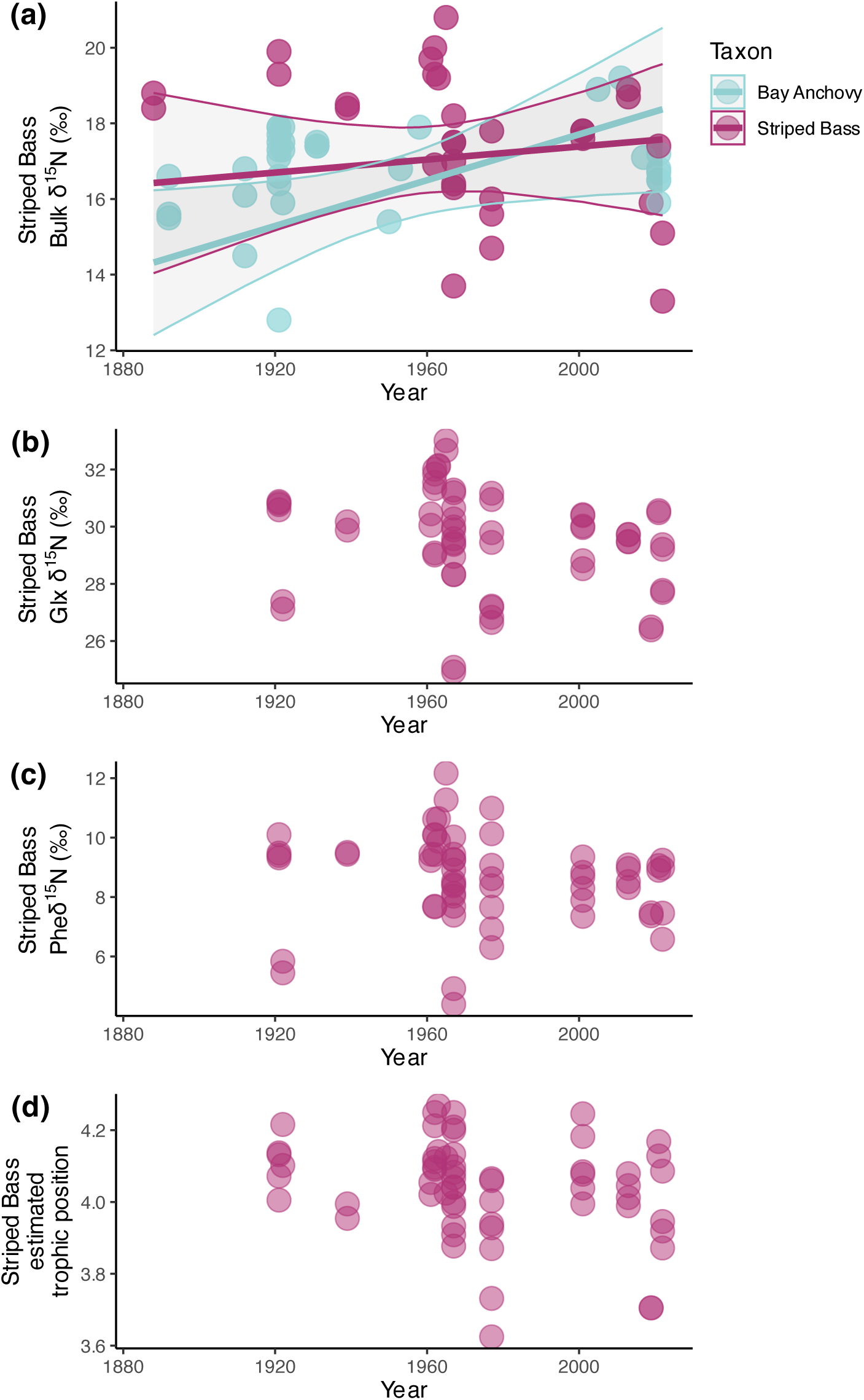
(a) Bulk (for all nitrogen atoms in the tissue sample, across different compounds) δ^15^N values for tissue samples from Striped Bass and Bay Anchovy. The axes variables are the same as in Figure 4a-e and the points for Striped Bass and Bay Anchovy are as shown in Figures 4a and d, included here to facilitate their comparison. The x-axis shows the year of specimen collection. Individual points show raw data. Fitted lines and ribbons represent predicted fitted values and their 95% credible intervals from the Bayesian model described in Table 1, for Striped Bass and Bay Anchovy (darker red-violet line representing fitted values for Striped Bass, matching the color of the points for Striped Bass as shown in the key, and the lighter blue-green colored line showing fitted values for Bay Anchovy). (b-d) Dataset analyzed by the model in Table 3: compound-specific δ^15^N values for Striped Bass specimens only. For each of the specimens, the results of two separate technical replicates for obtaining values of the y-axis variable are shown so that each specimen is plotted twice and represented by two replicate data points (60 points total). (d) Estimated trophic level (where trophic level equal to 1 represents the primary producers of a food web), which is calculated based on data shown in the two foregoing panels: (b) δ^15^N values in glutamic acid, and (c) δ^15^N values in phenylalanine. Estimated trophic level (d), δ^15^N values in glutamic acid (b), and δ^15^N values in phenylalanine (c) all have weak negative correlations with time (statistically insignificant in our modeling).

**Table 1.** Bulk (for all nitrogen atoms in the tissue sample, across different compounds) δ^15^N values of nitrogen (relative to amount of the nitrogen-15 isotope in atoms in N_2_ gas in air) for 183 tissue samples from five sampled Chesapeake Bay fish species (Striped Bass, Summer Flounder, Bluefish, Bay Anchovy, and Menhaden). δ^15^N value was analyzed as a Gaussian outcome with identity link in a generalized linear mixed-effects model fit with Bayesian model inference in brms. The 95% probability density credible intervals that do not overlap zero are highlighted in **bold**.

| Explanatory variable | Effect | Lower bound of 95% credible interval | Upper bound of 95% credible interval |
| --- | --- | --- | --- |
| Standard length (scaled) | **0.44** | **0.10** | **0.78** |
| Year (scaled) | **1.35** | **0.17** | **2.52** |
| Fresh and unpreserved? | -1.47 | -3.11 | 0.14 |
| Formalin-exposed? | -0.65 | -1.38 | 0.07 |
| **Taxon (offsets relative to Bay Anchovy, for the same given standard length)** |  |  |  |
| Menhaden | **-2.46** | **-3.24** | **-1.70** |
| Striped Bass | -0.18 | -1.39 | 1.02 |
| Summer Flounder | **-1.27** | **-2.24** | **-0.29** |
| Bluefish | 0.72 | -0.29 | 1.74 |
| **Watershed/region (offsets relative to mean for fishes from in/around Delmarva peninsula)** |  |  |  |
| James | -0.21 | -1.10 | 0.75 |
| York | -0.97 | -2.13 | 0.17 |
| Rappahannock | -0.05 | -1.28 | 1.17 |
| Potomac | 1.33 | -0.03 | 2.73 |
| Patuxent | **2.04** | **1.00** | **3.09** |
| Others between Patuxent and Susquehanna (inc. Susquehanna) | **1.67** | **0.94** | **2.41** |
| **Interactions (taxon effects are relative to Bay Anchovy)** |  |  |  |
| Year * Menhaden | **-0.94** | **-1.69** | **-0.18** |
| Year * Striped Bass | **-1.02** | **-1.92** | **-0.14** |
| Year * Summer Flounder | -0.38 | -1.31 | 0.56 |
| Year * Bluefish | **-0.93** | **-1.79** | **-0.08** |
| Year * James | -0.20 | -1.06 | 0.68 |
| Year * York | -0.14 | -1.28 | 0.98 |
| Year * Rappahannock | -0.06 | -1.21 | 1.06 |
| Year * Potomac | 0.06 | -1.99 | 2.12 |
| Year * Patuxent | 0.80 | -1.84 | 3.39 |
| Year * Other W Upper Chesapeake Bay | -0.04 | -1.07 | 1.00 |

Our data show effects on δ^15^N values in fish muscle of the river watersheds from which specimens were collected, but do not support potential differences between sites in how δ^15^N values varied through time. Samples from in and near the Patuxent River mouth and Maryland western shore showed δ^15^N values 2.81‰ points higher than samples from in and near the York River (Table 1). Interactions between watershed and year were not significant, with their credible intervals all broadly overlapping zero (Table 1). Supplemental Table 2 shows a version of the statistical model in Table 1 with additional covariates for Salinity and Season, neither of which shows credible interval estimates excluding zero. Supplemental Figure 2 shows the nonlinear nonparametric trends that were fit by a model using splines for river watershed-by-time relationships. As with nonlinear species-by-time interactions, including these splines resulted in a worse (higher) WAIC score for the statistical model, and does not qualitatively suggest that there are any nonlinear trends or differences between watersheds that were not captured with our linear models.

Across different years, there are consistent species-level differences in δ^15^N of individual fishes of a given standard length. Supplemental Figure 3 shows the variation across fishes’ standard lengths in δ^15^N values – standard length across all five species is estimated by our statistical model as having a significant positive effect on δ^15^N values (Table 1). The main effects of each taxon as shown in Table 1 are for fishes of a single, common standard length. The effect estimated for a specimen being of a given species, while holding standard length constant, is assuming standard length held constant at a single mean standard length value for the entire dataset across all five species. This common mean standard length value may or may not be a typical standard length value for that species. Consequently, the interpretation of the model saying that Bay Anchovy have on average slightly higher δ^15^N values than, for instance, Striped Bass in our data, is that they have higher δ^15^N after accounting statistically for the difference in body size between our Striped Bass specimens and our Bay Anchovy specimens, keeping in mind that the model is fitting a strong positive relationship between body size and δ^15^N. This aspect of the interpretation of the statistical model output means that the significant negative effect, for instance, of a specimen being a Summer Flounder as compared to a Bay Anchovy in its δ^15^N values (Table 1) applies for a Summer Flounder and Bay Anchovy at the same given standard length, and does not mean that a Summer Flounder of a much larger standard length than a Bay Anchovy would have a lower δ^15^N value than that Bay Anchovy. This case, of two individuals of the same standard length between these two species, does not occur in our dataset– our Summer Flounder specimens are all longer than our Bay Anchovy specimens. Nevertheless, our Bay Anchovy specimens may have similar values of δ^15^N, (see Supplemental Figure 3), as the much larger fishes from the three higher-trophic-level taxa in our study, and they have much higher δ^15^N values than Menhaden.

### Radiographs of gut contents: evidence to support assumptions about diets

Observations from radiographs support our assumption that small fishes comprised a significant part of the predator fish diets. We found that 52% of the X-rayed Bluefish (N = 28 total fish X-rayed), 32% of the Striped Bass (N = 31), and 12% of Summer Flounder (N = 28) specimens have vertebrate remains in their guts. We used a mixed-effects binomial-likelihood generalized linear model including standard length as a covariate to evaluate whether presence of vertebrate remains in X-rayed guts varied with time. We did not find significant effects of year of collection on presence of vertebrate remains (Table 2). Bluefish show an estimated reduction in log odds of fish prey presence of −1.46 (95% CI −4.19 to +0.54, posterior probability density for the effect = 92.24%). The credible interval for this decline thus includes zero and is not significant. Bayesian probabilities of either positive or negative time trends in presence of these vertebrate remains in guts are estimated at <80% for Striped Bass and Summer Flounder. Supplemental Figure 4 shows an example image of evidence of vertebrate prey radiographed in a museum fish specimen’s gut.

**Table 2.** Estimated effects of time (year of collection) and standard length on the log of the odds ratio of presence or absence of vertebrate prey in the guts of a collection specimen, from a Binomial generalized linear mixed-effects model. Higher coefficients mean higher odds of presence of vertebrate prey recorded with higher values of that explanatory variable. The model output in the table below is for three models run separately with the specimens from each of the three higher-trophic-level species in our study: 28 Bluefish, 31 Striped Bass, and 28 Summer Flounder.

| **Explanatory variable** | **Effect** | **Lower bound of 95% credible interval** | **Upper bound of 95% credible interval** |
| --- | --- | --- | --- |
| **Bluefish** |  |  |  |
| **Standard length (scaled)** | **-3.01** | **-6.98** | **-0.24** |
| Year (scaled) | -1.46 | -4.19 | 0.54 |
| **Striped Bass** |  |  |  |
| Standard length (scaled) | -1.30 | -5.06 | 0.77 |
| Year (scaled) | 1.17 | -1.78 | 6.39 |
| **Summer Flounder** |  |  |  |
| Standard length (scaled) | 1.75 | -1.21 | 5.27 |
| Year (scaled) | 0.29 | -3.35 | 4.08 |

### Compound-specific δ^15^N analysis: directly estimating trophic levels

Compound-specific isotope data do not support the hypothesis that Striped Bass in the Chesapeake Bay are changing in trophic level relative to Bay Anchovy, nor do they significantly support the suggestion from the bulk isotope data of a change in δ^15^N values at the base of the food web affecting all species. The estimated trophic level of Striped Bass does not show a significant trend through time (Table 3, Figure 5D). Estimated trophic level of the Striped Bass is negatively correlated with time, but not with a 95% credible interval excluding 0. Additionally, δ^15^N value of phenylalanine does not vary significantly through time (Figure 5C). Accounting for season covariates (Spring, Summer, Fall, or Winter, see Supplemental Table 3) did not affect model results. The variation in estimated trophic level between seasons is shown in Supplemental Figure 5, but such variation was not significant (Supplemental Table 3). The values of the estimated trophic levels ranged from about 3.6 for a 24 cm SL fish to about 4.3 for a 36 cm SL fish – there was a slight correlation (linear Pearson’s correlation of 0.13) between SL and estimated trophic level in the data, but the estimated positive effect of SL on trophic level in these data was not significant in the model in Table 3. A lack of consistent, significant variation in estimated trophic level with time and other covariates implies that glutamic acid δ^15^N and phenylalanine δ^15^N must closely correlate. They must do so because glutamic acid δ^15^N is affected more by trophic level whereas phenylalanine δ^15^N is not as affected (Chikaraishi et al. 2009). And, the difference between δ^15^N in these two amino acids corresponds to the estimated trophic level according to the calculation shown in the Methods. Therefore, if the estimated trophic level (proportional to the distance in δ^15^N between these two compounds) is not changing, the δ^15^N of these two amino acids must not be varying through time relative to each other. These amino acids’ δ^15^N values indeed do closely correlate with each other, and with the bulk δ^15^N values for fish that were part of both our CSIA data collection and our bulk isotopes analyses (Supplemental Figure 6).

**Table 3.** Estimated trophic level of 30 Striped Bass specimens, fit as a Gaussian outcome with identity link in a generalized linear mixed-effects model fit with Bayesian model inference in brms.

| **Explanatory variable** | **Effect** | **Lower bound of 95% credible interval** | **Upper bound of 95% credible interval** |
| --- | --- | --- | --- |
| Standard length (scaled) | 0.03 | -0.02 | 0.08 |
| Year (scaled) | -0.04 | -0.09 | 0.01 |

## Discussion

For Chesapeake Bay Striped Bass, a dominant predator in the largest estuary in North America, we find no evidence for shifts over time in trophic level. Compound-specific isotope analyses do not show a significant directional change through time in estimated trophic level. The Striped Bass in our study vary around a trophic level of 4 (tertiary consumer) across years and sites. This value is slightly lower than other trophic level estimates for this species from larger individuals. Feiner et al. (2013) compared Striped Bass bulk δ^15^N to those of nearby benthic invertebrates to normalize them and estimate trophic level and obtained a mean trophic level of 5.1 for 40-70 cm total length (TL) Striped Bass in North Carolina reservoirs in 2010. Even early in our time series, in the 1920s, estimated Striped Bass trophic level is near 4, so this low trophic level in our estimates does not reflect a decline over time. Disruptions to ecosystems from harvest and other modern industrialized human impacts can “flatten” food webs and reduce the trophic level of consumers by limiting prey choices (Dell et al. 2015). Despite this potential for “flattening”, some fishes show evidence of increasing trophic levels in the face of changing environments (Durante et al. 2022; Welicky et al. 2023). The diversity of prey items that Chesapeake Bay Striped Bass consume–over 220 different taxa recorded from gut metabarcoding (Pagenkopp Lohan et al. 2023)–may buffer them against trophic downgrading by reduced choice in prey. Trophic level of this taxon in this region may be staying constant over time even as multiple other characteristics of its trophic niche are changing (as observed in other systems, e.g., Everglades fishes in (Flood et al. 2023).

A stable trophic level through time for our Striped Bass is consistent with other studies of multiple fish species that have drawn on gut content data (Sánchez-Hernández et al. 2022) or isotopes (Nakazawa et al. 2010) to show that length-specific diets and ontogenetic diet shifts remain constant over decadal timescales. Other studies that used compound specific isotope analyses on fishes from archival collections, with similar sample sizes and statistical power to our study, have also found trophic level stasis through time. Sabadel et al. (2020) conducted compound-specific isotope analysis for specimens from 1955 to the present of three coastal New Zealand marine fishes and found significant increase through time in trophic level in the Tarakihi, *Nemadactylus macropterus* (from 19 specimens), but no change in two other taxa. Welicky et al. (2023) likewise found an increase through time in trophic level of one of five sampled Puget Sound marine fish species (50 specimens sampled/species), but no trends in the other four. Willert et al. (2023) used preserved primary producer (algae) specimens for information about food web baseline ^15^N (rather than compound-specific analysis), and estimated significant trophic level declines of two fishes over the last century, using 60 specimens per species. However, Willert et al.’s estimated effect sizes for trophic level decline were larger in magnitude than the weak negative correlation we see between our estimated Striped Bass trophic level and time.

Our bulk δ^15^N data are consistent with anthropogenic nitrogen input as a main driver of fish tissue isotopic variation in Chesapeake Bay. If anthropogenic nitrogen, which tends to increase δ^15^N values (Bucci et al. 2007; Baeta et al. 2017), was dominating isotopic variation in our study, we would expect to see δ^15^N value increases over time affecting both prey and predators, persisting after including covariates for seasonal effects. These seasonal effects can affect nitrogen dynamics by changing both freshwater inflows and mixing (Horrigan et al. 1990). We might expect that watersheds with consistently higher human population density and human impacts throughout the past two centuries would have higher fish muscle tissue δ^15^N values.

With anthropogenic nitrogen input rising throughout the Chesapeake Bay–not plateauing until at least the 1980s (Kemp et al. 2005)–we would not necessarily expect strong interactions between the time trend of δ^15^N values and our watershed covariate, as all watersheds would be increasing in δ^15^N values. Many of these expectations are supported in our dataset. The West Upper Chesapeake Bay and the Potomac River have since 1900 been highest in human population density as compared to other places in Chesapeake Bay (Paullin and Wright 1932), in ways that affected the δ^15^N values entering organisms in these watersheds. Black et al. (2017) showed more increase between 1750-1800 and 1850-1900, than between 1850-1900 and modern (2012-2013) samples, in δ^15^N values of historical oyster shells in the Edgewater, Maryland area in the Upper Chesapeake Bay, less than 50 miles North of the Patuxent River mouth. The much lower δ^15^N values in fishes in our study collected from the York watershed as compared to the Patuxent and Upper Chesapeake Bay watersheds is consistent with a role for an increased flow of anthropogenic nitrogen from developed areas. The expectation that is least supported by our data is that we would observe increases through time in δ^15^N values in fish body tissues. Our statistical models estimate this result for Bay Anchovy, but slightly less so for Summer Flounder, and much less so for Bluefish, Striped Bass, or Menhaden.

Patterns of environmental variation apart from anthropogenic nutrients reaching Chesapeake Bay may play a role in isotopic variation in our data, but none is well-supported or easily assessed by our dataset. The York River is closer to the Atlantic Ocean than is the Upper Chesapeake Bay, and higher salinity and more marine inputs could play a role in its difference from other watersheds in our data. However, environmental nitrogen following the Chesapeake Bay mainstem toward the ocean does not tend to have consistently lower or higher δ^15^N values with proximity to the ocean (Horrigan et al. 1990), so it is not clear that marine vs. freshwater character *per se* of environments in the Chesapeake Bay would drive δ^15^N values’ variation.

Also, salinity, which we can use as a proxy for proximity to the ocean and more marine nitrogen sources, does not significantly explain isotopic variation in our data. Even if salinity does not causally influence δ^15^N values, it tends to be confounded and negatively spatially correlated with land development and use in the Chesapeake Bay (Davias et al. 2014). This expectation of a negative relationship between salinity and δ^15^N therefore asks why we do not see a more negative relationship between salinity and δ^15^N in our data. Our watershed covariate absorbs the variation from this negative correlation between-watersheds, but this negative correlation could still exist within a watershed. Ultimately, anthropogenic nitrogen inputs increasing δ^15^N values are not consistently higher or lower upriver or downriver in Chesapeake Bay’s watersheds but peak near specific centers of pollution (Pennino et al. 2016). The increasing imperviousness of terrestrial environments around Chesapeake Bay means that water can flow and carry pollutants long distances in unpredictable floods with little of the natural trapping or filtering of water. (Kemp et al. 2005).

Given the literature and data about modern nitrogen inputs and characteristics in aquatic ecosystems, it is surprising that Striped Bass do not show an increase through time in their muscle tissue δ^15^N values while their trophic level remained constant. Estuarine and coastal environments worldwide are increasingly flooded with anthropogenic nitrogen that increases organisms’ δ^15^N (Baeta et al. 2017) and we see some signal of this anthropogenic nitrogen enrichment in our study, mostly in our Bay Anchovy data. The closer correspondence between Bay Anchovy increase through time in δ^15^N values and the time trends observed in Summer Flounder δ^15^N values, as compared to Striped Bass and Bluefish δ^15^N values, is not surprising: across sites in Chesapeake Bay, Summer Flounder δ^15^N values closely track Bay Anchovy δ^15^N values and δ^15^N values follow similar seasonal changes between these two species (Buchheister and Latour 2011). Buchheister and Latour (2011) suggest this similarity and coupling is due to their similar use of benthic invertebrate prey items. As in our study, they found that Bay Anchovy may have bulk muscle tissue δ^15^N values of 18‰ or higher, similar to or higher than those of Summer Flounder in the same environment, even though Summer Flounder are generally much larger than Bay Anchovy and are Bay Anchovy predators. It is unclear why Striped Bass would not also show a pattern of increasing δ^15^N values with time, if such increases occur in Bay Anchovy and Summer Flounder due to increased anthropogenic nitrogen. Bay Anchovy and Summer Flounder are sometimes noted as being more dependent on deep-water, sub-pycnocline, and/or benthic habitats as compared to Bluefish, Striped Bass, and Menhaden (Batiuk et al. 2009), but Bay Anchovy are the single most common finfish prey of <50 cm TL Striped Bass in Chesapeake Bay (Overton et al. 2009) and are distributed in many of the same environments in Chesapeake Bay, and subject to similar limiting effects on their distribution by low oxygen in Chesapeake Bay’s shallow waters (Batiuk et al. 2009). As Menhaden populations have generally declined (though with substantial fluctuation) since the 1950s (Love et al. 2006), gut content evidence suggests that Striped Bass are eating proportionally more Bay Anchovy and benthic invertebrates (Overton et al. 2015) and more benthic fishes (Walter and Austin 2003) compared to previous dominance of Menhaden in the 1950s Chesapeake Bay Striped Bass diet (Griffin and Margraf 2003). ^13^C isotopes also indicate they are eating proportionally more benthic invertebrates (Pruell et al. 2003) – diet shifts that could cause them to be more, not less, likely to mirror δ^15^N increases seen in benthic invertebrate-eating Bay Anchovy and Summer Flounder.

The Striped Bass in our study do not show evidence of being in a food web where δ^15^N is increasing in nitrogen sources at the base of that food web. It is not surprising, given their diverse diet and potential slight habitat differences with Bay Anchovy, that Striped Bass do not perfectly mirror Bay Anchovy isotope trends. But it is unclear what is contributing to δ^15^N in their other prey (much of which is neither Bay Anchovy or Menhaden) that leads to Striped Bass δ^15^N values not increasing over time. In ecosystems where increased anthropogenic nitrogen leads to δ^15^N enrichment, such trends often affect fish species across the entire community, not just certain fish species to the exclusion of others (e.g., Turner et al. 2015). However, phenylalanine δ^15^N values in Striped Bass, which should reflect variation in δ^15^N at the base of its food web (Chikaraishi et al. 2009), do not show any increase through time. Although we can provide possible explanations for this discrepancy between the compound-specific and bulk isotope data, we cannot resolve this discrepancy with our dataset. Even though Striped Bass diets in Chesapeake Bay are sometimes over 50% Bay Anchovy by weight (Overton et al. 2009), we should not expect Bay Anchovy δ^15^N values trends to perfectly predict those of Striped Bass.

But, why the other prey species of Striped Bass are not passing to it these same inflated δ^15^N values seen in Bay Anchovy (and filter-feeding oysters, Black et al. 2017) is unclear and speaks to the complexity of food webs in Chesapeake Bay. Thus, there is not a single “base” to the food web in Chesapeake Bay from which bottom-up nutrient dynamics flow. Striped Bass have seasonal variation in diet throughout Chesapeake Bay, with invertebrates increasing in Striped Bass diet in the Spring in all regions (Overton et al. 2009). But our model fitting did not indicate Season as a significant driver of variation in our Striped Bass data, and including such patterns did not change Year effects.

One factor likely influencing the nitrogen isotope contents of Striped Bass tissue in our study and driving variation between individuals is that Chesapeake Bay Striped Bass are anadromous and may spend up to the first two or three years of their life far upriver in freshwater or near-fresh water (Gahagan et al. 2015). Small Striped Bass of the standard lengths in our study, even if collected in Chesapeake Bay itself, may only weeks or months earlier been further upriver. Although little is known about how quickly the signal of isotope ratios from particular diet items turn over in Striped Bass as an individual grows and its diet changes (Baker et al. 2016), a recent migrant’s isotope ratios may reflect more about where they lived (and what they ate there) than it does about their diet in their new habitat. Substantial variation exists between individual Striped Bass in their migratory behavior (Secor et al. 2020), likely adding inter-individual variation to our data. We do not know the migratory histories of the museum specimens in our data, so the influence of this movement is unknown. Young Chesapeake Bay Striped Bass are affected differently in nitrogen isotope ratios by freshwater as compared to more marine environments (Conroy et al. 2015). This difference may be a factor in differences between temporal trends in Striped Bass in our data and temporal trends in the other four species in our data that did not inhabit the same early-life environments. However, δ^15^N values in more freshwater Striped Bass are slightly higher than those in more brackish individuals (Conroy et al. 2015). Freshwater environments for Striped Bass may have different time trends in environmental nitrogen sources, diet items available for young Striped Bass, or both; whether these trends differ is not known and there is not any evidence that freshwater δ^15^N values in recent years would systematically depress Striped Bass tissue δ^15^N values relative to other fishes.

Bay Anchovy we measured for this study were mostly from different sites than our Striped Bass and Bluefish. We may, therefore, question how representative our Bay Anchovy specimens are of the Bay Anchovy these predators consume, although all of these species are found at and can move to all collection sites. One consideration is how high the bulk isotope values are in both our and Buchheister and Latour’s study for Bay Anchovy, and the strong length-specific variation in δ^15^N values of Bay Anchovy we observe. Our data (Supplemental Figure 3) and other work suggest the ability of Bay Anchovy ability to acquire certain prey (Houde and Zastrow 1991) and avoid certain predators is closely tied to measures of their body length. Larger-bodied Bay Anchovy may be rarer in the modern Chesapeake Bay (Jung and Houde 2004), although more data is needed to confirm this possibility. Striped Bass and Bluefish consuming particularly small Bay Anchovy would not increase their δ^15^N values as much as if they were eating large Bay Anchovy. The net energetic profitability of Bay Anchovy as prey items, for Bluefish that are similar to lengths considered in our study, declines with Bay Anchovy length (Scharf et al. 2002). Therefore, relatively large Bay Anchovy may not have been pursued by Bluefish in our study. If our Striped Bass were eating disproportionately more lower-trophic-level, small-bodied Bay Anchovy, this increased consumption of small fish could reduce their trophic level. The rest of their diet must have maintained their trophic level, given that our compound-specific results do not show a significant trophic level decline.

Reconstructing diets of fishes from indirect lines of evidence is challenging, especially for fishes in past time periods. Yet such knowledge is valuable for ecosystem-based conservation and management (Buchheister et al. 2015) Our study and its limitations illustrate key insights. Our study speaks to the importance of integrating multiple sources of data on the length-specific diet of a population before drawing conclusions about patterns of change in ecological interactions. Our study is consistent with experimental findings that time under exposure to preservatives does not significantly alter the size of preservation effects in fishes’ δ^15^N values (Durante et al. 2020). However, similar to previous similar preservation experiments, our experiment was not on the timescale of over a hundred years on which these fish have been collected and stored, meaning our understanding of preservation effects in our data over decades may be incomplete. Our study also shows the value of specimens of both predatory species and their prey species collected from the same localities and times at a fine scale to be able to compare data from a predator with specimens that represent prey from the same habitat.

Specimens of predators and their prey–or other ecological interactors, like host and parasite–collected in situ from the same time and place, are valuable in historical ecology (Haug et al. 2023; Wood et al. 2023). Collection of more such specimens is therefore valuable. Our study also highlights the potential value of museum collections of taxa sampled across space and time, as archives of ecological information and environmental change. Museum specimens are even more valuable when more data are available about their environmental context (Rocha et al. 2014). Lastly, our work highlights how fruitful it is to assemble knowledge of a fish’s prey items, to collect data on not just the species a predator consumes but in which environments they consume that prey and which ages and body sizes of that prey species a given predator prefers or is able to consume (Hartman 2000). These are all important insights with which to approach the complex task of characterizing variation in the diets of the fishes of Chesapeake Bay and beyond.

## Acknowledgments

Dr. Christine France and the Stable Isotope Facility at the Smithsonian Institution Museum Conservation Institute performed bulk isotope analysis after the first author prepared samples for spectrometry. The UC Davis Stable Isotope Facility performed compound-specific isotope analysis. Drs. Mary Fabrizio, Troy Tuckey, and Wendy Lowery secured newly collected fish specimens from Virginia Institute of Marine Science (VIMS) survey work for this study. Drs. Eric Hilton and Sarah Huber, and Miguel Montalvo, assisted with use of VIMS fluid-preserved collections. Diane Pitassy, LaShaun Willis, Dr. Abby Reft, and Ella Haigler assisted with use of the Smithsonian Institution National Museum of Natural History fish collection, Museum Support Center lab spaces, and X-ray procedures. Drs. Joe Travis, Mariana Fuentes, Kimberly Hughes, and Michael Cortez provided feedback on the design of this project and comments on revisions of this manuscript. Two anonymous reviewers provided extensive, helpful feedback on an earlier version of this paper. Silhouettes in Figures 1 and 4, and in Supplemental Figure 3, are from PhyloPic.org under a CC0 1.0 Universal Public Domain Dedication license (no permission or attribution required); these images were created by Rene Martin (large fish), Matt Kolmann (“smaller fish of trophic level =∼3”), Nathan Hermann (other small fish), Guillaume Dera (blue crab, dinoflagellate), Nathan Jay Baker (amphipod), Rachel Wooliver (seagrass), Levi Simons (green algae), and Sarah Frail (diatoms) (the original art was modified to have a consistent solid black color for all animal silhouettes).

## Declarations

### Ethics

No ethics approvals were required for this work.

### Consent to participate

Not applicable – no human subjects.

### Availability of data and material

All isotope data and a list of Specimens Examined are included as Electronic Supplementary Material.

### Code availability

A code file with scripts used for statistical output tables in this manuscript is included as Electronic Supplementary Material.

