## Supplemental Figures and Tables for "Estimating Historical Food Web Variation in Chesapeake Bay Using Isotope Variation in Museum Fish Specimens"

### SUPPLEMENT

Supplemental Table 1. Total numbers of specimens sampled and included in the statistical analyses (shown in Table 1) for different combinations of locality, time period, and group of species. “HTL” refers to higher trophic level taxa Striped Bass, Summer Flounder, and Bluefish; “LTL” refers to lower trophic level taxa Bay Anchovy and Menhaden. For each group of specimens represented by each interior cell, the number of specimens (n) is given, as well as the mean standard length, body weight, and mean year-averaged salinity ($\mu_L$, $\mu_W$, and $\mu_S$) for each group of specimens.

| Species | 1850-1900 | 1900-1950 | 1950-2000 | 2000-2022 |
|---|---|---|---|---|
| | Closest to **James** River mouth and watershed | | | |
| HTL (higher trophic level) taxa, Apr-Sep | n=5, $\mu_L$=22.2, $\mu_W$=157.9, $\mu_S$=18 | n=5, $\mu_L$=17, $\mu_W$=78, $\mu_S$=21 | n=0 | n=1, $\mu_L$=17, $\mu_W$=115.6, $\mu_S$=24 |
| HTL taxa, Oct-Mar | n=1, $\mu_L$=22, $\mu_W$=147.9, $\mu_S$=18 | n=0 | n=0 | n=2 (fresh), $\mu_L$=32, $\mu_W$=653, $\mu_S$=18 |
| LTL (lower trophic level) taxa, Apr-Sep | n=4, $\mu_L$=5.6, $\mu_W$=1.7, $\mu_S$=18 | n=1, $\mu_L$=13, $\mu_W$=30.4, $\mu_S$=18 | n=1, $\mu_L$=15, $\mu_W$=94.2, $\mu_S$=0.5 | n=0 |
| LTL taxa, Oct-Mar | n=4, $\mu_L$=23.8, $\mu_W$=191.9, $\mu_S$=18 | n=1, $\mu_L$=6.8, $\mu_W$=2.9, $\mu_S$=21 | n=0 | n=0 |
| | Closest to **York** or **Rappahannock** River mouth and watershed | | | |
| HTL taxa, Apr-Sep | n=0 | n=0 | n=5, $\mu_L$=19.6, $\mu_W$=119.8, $\mu_S$=11 | n=4, $\mu_L$=24.1, $\mu_W$=228.5, $\mu_S$=5.6 |
| HTL taxa, Oct-Mar | n=0 | n=6, $\mu_L$=12.8, $\mu_W$=26.3, $\mu_S$=18 | n=7, $\mu_L$=18.6, $\mu_W$=102.5, $\mu_S$=7.1 | n=8 (fresh), $\mu_L$=26.8, $\mu_W$=414.8, $\mu_S$=12.1 |
| LTL taxa, Apr-Sep | n=2, $\mu_L$=8.5, $\mu_W$=10.8, $\mu_S$=9 | n=0 | n=0 | n=2, $\mu_L$=12.5, $\mu_W$=59.7, $\mu_S$=10 |
| LTL taxa, Oct-Mar | n=0 | n=0 | n=0 | n=12 (11 fresh), $\mu_L$=9.6, $\mu_W$=27.7, $\mu_S$=14.2 |
| | Closest to **Potomac** River mouth and watershed | | | |

| | | | | |
|---|---|---|---|---|
| HTL taxa, Apr-Sep | n=0 | n=11, $\mu_L$=19.7, $\mu_W$=179.3, $\mu_S$=8.4 | n=2, $\mu_L$=39, $\mu_W$=1088.1, $\mu_S$=12.5 | n=0 |
| HTL taxa, Oct-Mar | n=0 | n=0 | n=0 | n=0 |
| LTL taxa, Apr-Sep | n=2, $\mu_L$=11, $\mu_W$=18.8, $\mu_S$=10 | n=6, $\mu_L$=10, $\mu_W$=31.4, $\mu_S$=7.9 | n=0 | n=0 |
| LTL taxa, Oct-Mar | n=0 | n=6, $\mu_L$=6.3, $\mu_W$=4.4, $\mu_S$=8.8 | n=0 | n=0 |
| | Closest to **Patuxent** River mouth and watershed, to **Susquehanna**, or other W. Upper Chesapeake Bay | | | |
| HTL taxa, Apr-Sep | n=2, $\mu_L$=34.5, $\mu_W$=646.1, $\mu_S$=0 | n=12, $\mu_L$=14.6, $\mu_W$=48.7, $\mu_S$=10.4 | n=6, $\mu_L$=20.7, $\mu_W$=235.6, $\mu_S$=8.3 | n=5, $\mu_L$=15.4, $\mu_W$=52.4, $\mu_S$=8.5 |
| HTL taxa, Oct-Mar | n=0 | n=2, $\mu_L$=13, $\mu_W$=35.1, $\mu_S$=7.5 | n=2, $\mu_L$=21.8, $\mu_W$=140.5, $\mu_S$=11.3 | n=0 |
| LTL taxa, Apr-Sep | n=0 | n=15, $\mu_L$=6.8, $\mu_W$=5.7, $\mu_S$=7.8 | n=4, $\mu_L$=8, $\mu_W$=7, $\mu_S$=7 | n=2, $\mu_L$=10, $\mu_W$=15.7, $\mu_S$=7.5 |
| LTL taxa, Oct-Mar | n=0 | n=4, $\mu_L$=12.5, $\mu_W$=35, $\mu_S$=8.1 | n=0 | n=3, $\mu_L$=9, $\mu_W$=15.3, $\mu_S$=7.5 |
| | Delmarva | | | |
| HTL taxa, Apr-Sep | n=0 | n=6, $\mu_L$=20.6, $\mu_W$=156.7, $\mu_S$=17 | n=3, $\mu_L$=20.2, $\mu_W$=161.5, $\mu_S$=26.7 | n=0 |
| HTL taxa, Oct-Mar | n=0 | n=7, $\mu_L$=13.6, $\mu_W$=41.2, $\mu_S$=21 | n=1, $\mu_L$=30.5, $\mu_W$=393.6, $\mu_S$=10 | n=5 (fresh), $\mu_L$=21.4, $\mu_W$=173.8, $\mu_S$=17.4 |
| LTL taxa, Apr-Sep | n=0 | n=1, $\mu_L$=6, $\mu_W$=3.1, $\mu_S$=15 | n=2, $\mu_L$=4.9, $\mu_W$=1.5, $\mu_S$=15 | n=0 |
| LTL taxa, Oct-Mar | n=0 | n=2, $\mu_L$=13.5, $\mu_W$=46.5, $\mu_S$=12.5 | n=1, $\mu_L$=6, $\mu_W$=1.9, $\mu_S$=7.5 | n=0 |

Supplemental Table 2. Bulk (for all nitrogen atoms in the tissue sample, across different compounds) $\delta^{15}N$ values of nitrogen (relative to amount of the nitrogen-15 isotope in atoms in $N_2$ gas in air) for 183 tissue samples from five sampled Chesapeake Bay fish species (Striped Bass, Summer Flounder, Bluefish, Bay Anchovy, and Menhaden), with additional covariates. $\delta^{15}N$ value was analyzed as a Gaussian outcome with identity link in a generalized linear mixed-effects model fit with Bayesian model inference in brms. The 95% probability density credible intervals that do not overlap zero are highlighted in **bold**. Additional covariates, Salinity and Season, are added in this model as compared with Table 1.

| **Explanatory variable** | **Effect** | **Lower bound of 95% credible interval** | **Upper bound of 95% credible interval** |
|---|---|---|---|
| Standard length (scaled) | **0.41** | **0.07** | **0.76** |
| Year (scaled) | **1.33** | **0.17** | **2.50** |
| Fresh and unpreserved? | **-1.78** | **-3.43** | **-0.15** |
| Formalin-exposed? | 0.51 | -0.26 | 1.27 |
| Salinity | -0.11 | -0.42 | 0.22 |
| Season (Winter) | 0.58 | -0.01 | 1.19 |
| **Taxon (offsets relative to Bay Anchovy, for the same given standard length)** | | | |
| Menhaden | **-2.41** | **-3.20** | **-1.63** |
| Striped Bass | -0.11 | -1.31 | 1.11 |
| Summer Flounder | -1.00 | -2.02 | 0.02 |
| Bluefish | 0.84 | -0.15 | 1.84 |
| **Watershed/region (offsets relative to mean for fishes from in/around Delmarva peninsula)** | | | |
| James | -0.08 | -1.05 | 0.89 |
| York | -0.94 | -2.12 | 0.20 |
| Rappahannock | -0.16 | -1.49 | 1.16 |
| Potomac | **1.47** | **0.07** | **2.89** |
| Patuxent | **1.96** | **0.86** | **3.05** |
| Others between Patuxent and Susquehanna (inc. Susquehanna) | **1.64** | **0.78** | **2.51** |
| **Interactions (taxon effects are relative to Bay Anchovy)** | | | |
| Year * Menhaden | **-0.87** | **-1.63** | **-0.11** |
| Year * Striped Bass | **-0.91** | **-1.80** | **-0.02** |
| Year * Summer Flounder | -0.37 | -1.27 | 0.55 |

| | | | |
|---|---|---|---|
| Year * Bluefish | -0.81 | -1.67 | 0.04 |
| Year * James | -0.24 | -1.11 | 0.63 |
| Year * York | -0.16 | -1.26 | 0.96 |
| Year * Rappahannock | -0.08 | -1.25 | 1.09 |
| Year * Potomac | 0.27 | -1.80 | 2.36 |
| Year * Patuxent | 0.51 | -2.12 | 3.12 |
| Year * Other W Upper Chesapeake Bay | -0.15 | -1.20 | 0.90 |

Supplemental Table 3. Estimated trophic level of 30 Striped Bass specimens, fit as a Gaussian outcome with identity link in a generalized linear mixed-effects model fit with Bayesian model inference in brms. As compared with Table 3, this model includes Season covariates.

| **Explanatory variable** | **Effect** | **Lower bound of 95% credible interval** | **Upper bound of 95% credible interval** |
|---|---|---|---|
| Standard length (scaled) | 0.04 | -0.02 | 0.10 |
| Year (scaled) | -0.04 | -0.09 | 0.02 |
| Season – Summer (relative to Spring) | -0.06 | -0.20 | 0.08 |
| Season - Fall | 0.03 | -0.09 | 0.16 |
| Season - Winter | -0.03 | -0.23 | 0.16 |

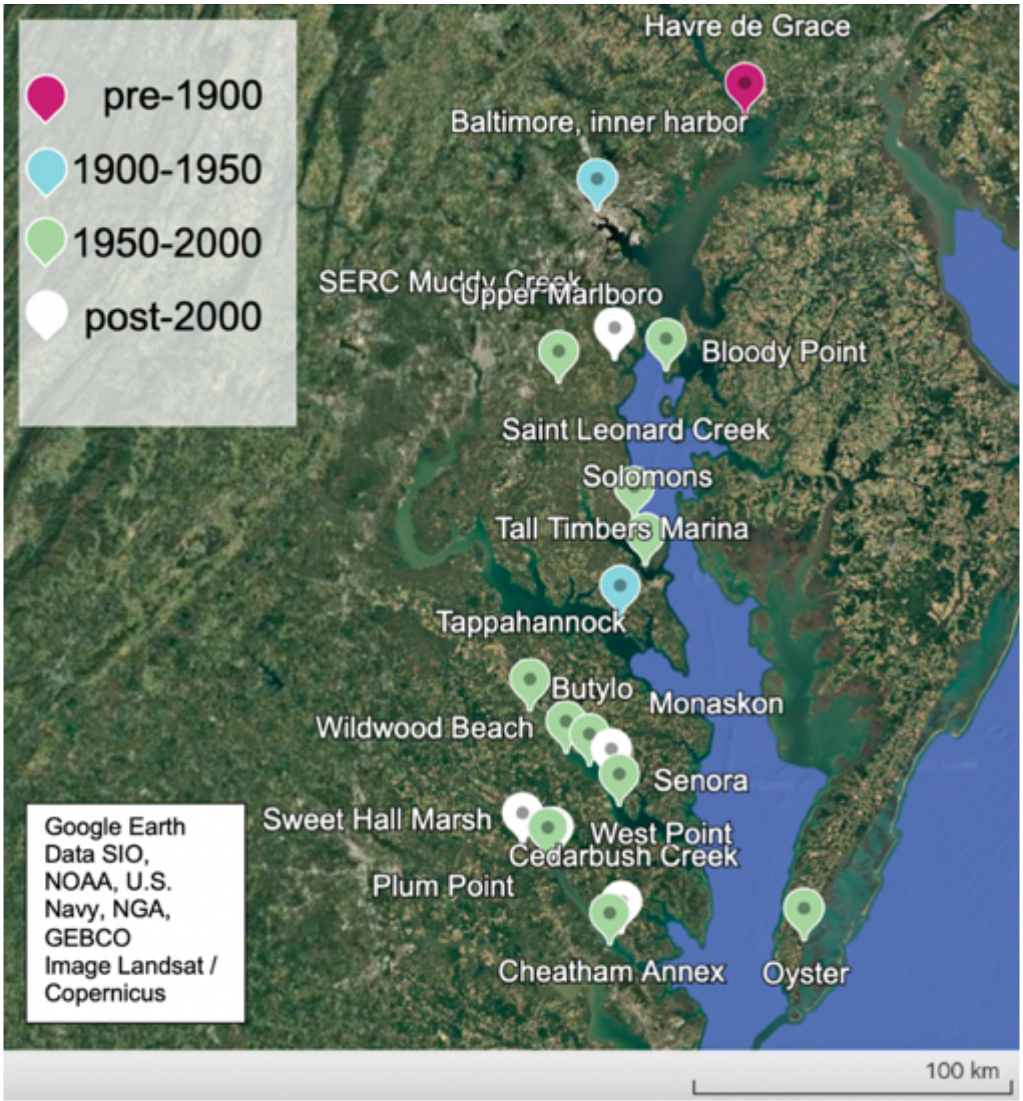


Supplemental Figure 1. A Google Earth map of the collection site and time period of the Striped Bass specimens used in this study. Map data: Google Landsat / Copernicus, SIO, NOAA, U.S. Navy, NGA, GEBCO. All sites referenced in the Supplemental Data Table for the entire study can be found at this web-based map: https://earth.google.com/earth/d/1VIAwCos29QDZRvoz10ZMTPySRsGCW6wl?usp=sharing.

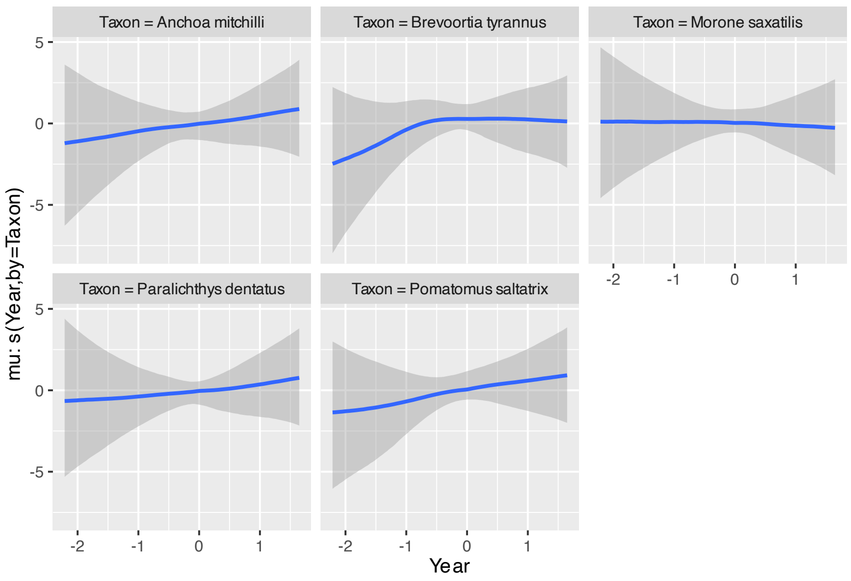


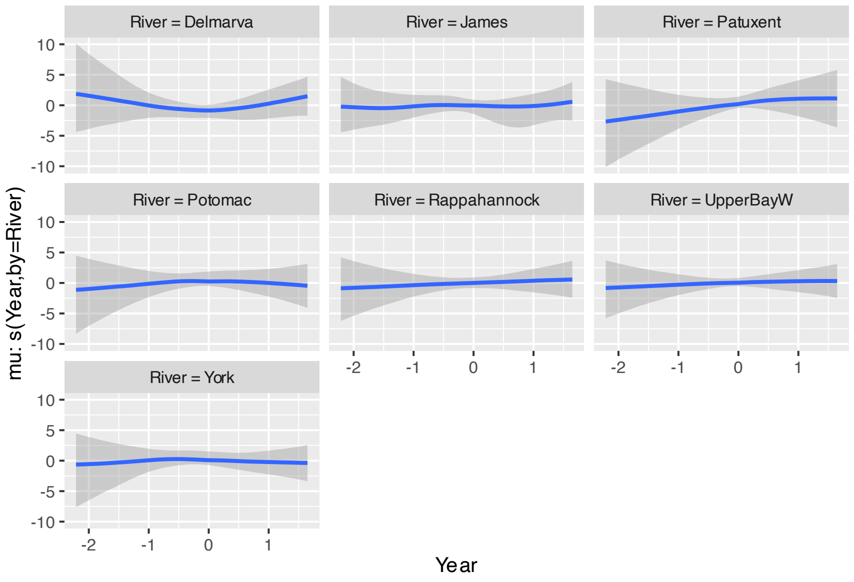


Supplemental Figure 2. Nonlinear nonparametric spline fits for variation in $\delta^{15}N$ values (on the y-axis) across years included in our study (on the x-axis), broken down by different watersheds and taxa.

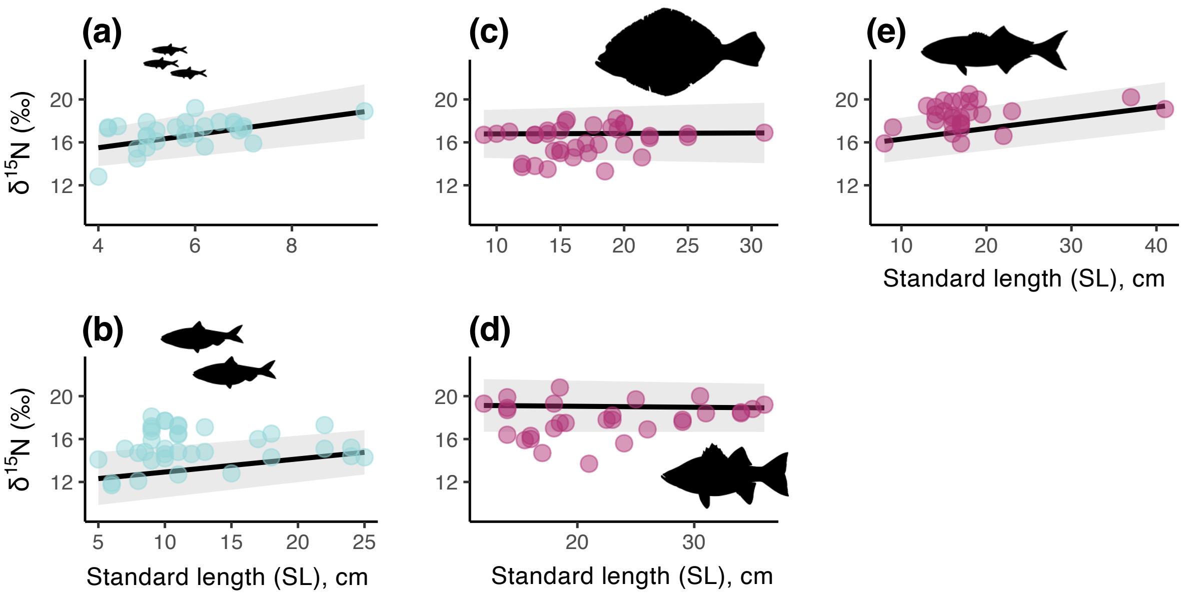


Supplemental Figure 3. (a-e) Bulk (for all nitrogen atoms in the tissue sample, across different compounds) $\delta^{15}N$ values (relative to amount of the nitrogen-15 isotope in atoms in $N_2$ gas in air) for 157 preserved museum specimen tissue samples from five sampled Chesapeake Bay fish species. (a) Bay Anchovy, (b) Menhaden, (c) Summer Flounder, (d) Striped Bass, and (e) Bluefish. The x-axes show the standard length of the specimen. Individual points show raw data. The fitted line and confidence interval are for an exploratory LOESS smoother evaluated at two points, to show the general linear trend in the data prior to addition of any other explanatory covariates.

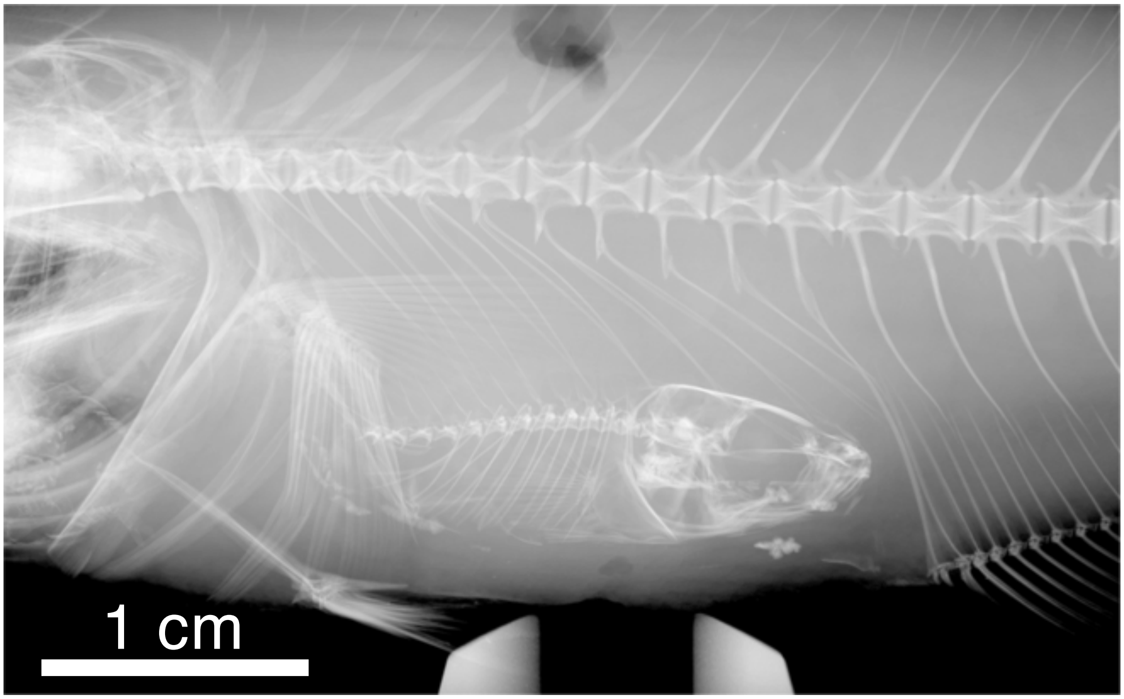


Supplemental Figure 4. X-ray of Bluefish, USNM 89800, with fish prey in gut, collected August 1930, Cobb Island, Maryland (Potomac River).

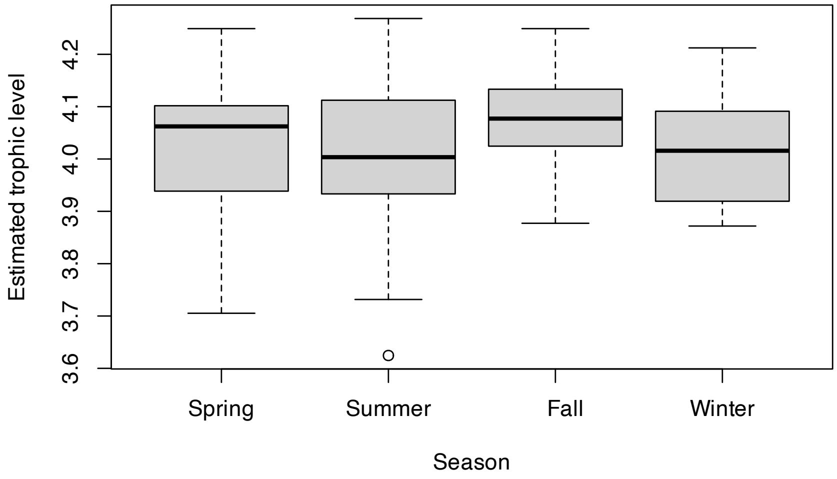


Supplemental Figure 5. Variation in Striped Bass estimated trophic levels across seasons of specimen collection.

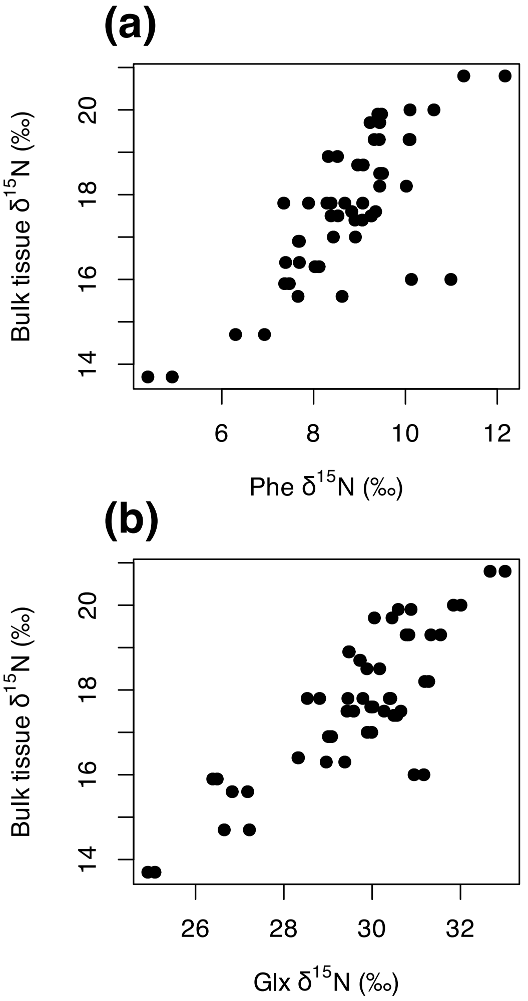


Supplemental Figure 6. For the 30 Striped Bass specimens in Figure 5b-d, (a) the $\delta^{15}N$ values in phenylalanine plotted on the x-axis and the samples' corresponding bulk muscle tissue $\delta^{15}N$ values plotted on the y-axis. (b) the $\delta^{15}N$ values in glutamic acid plotted on the x-axis and the samples' corresponding bulk muscle tissue $\delta^{15}N$ values plotted on the y-axis.
